# *KRAS* allelic imbalance drives tumour initiation yet suppresses metastasis in colorectal cancer *in vivo*

**DOI:** 10.1101/2023.08.29.555396

**Authors:** Arafath K. Najumudeen, Sigrid K. Fey, Laura M. Millett, Catriona Ford, Kathryn Gilroy, Nuray Gunduz, Rachel A. Ridgway, Eve Anderson, Douglas Strathdee, William Clark, Colin Nixon, Jennifer P. Morton, Andrew D. Campbell, Owen J. Sansom

## Abstract

Oncogenic *KRAS* mutations are well-described functionally and are known to drive tumorigenesis. Recent reports describe a significant prevalence of *KRAS* allelic imbalances or gene dosage changes in human cancers, including loss of the wild-type allele in *KRAS* mutant cancers. However, there is still much debate over the function of wild-type *KRAS* in tumour initiation, progression and therapeutic response. We have developed a genetically engineered mouse model which allows deletion of the wild-type copy of *Kras* in the context of an intact oncogenic *Kras* in colorectal cancer. We observe that in the presence of oncogenic *Kras*, wild-type *Kras* acts to restrain tumour growth. Mechanistically, deletion of wild-type *Kras* exacerbates oncogenic KRAS signalling through MAPK and thus drives tumour initiation. Absence of wild-type *Kras* potentiates the oncogenic effect of KRASG12D, while presence of wild-type *Kras* is associated with resistance to inhibition of MEK1/2 in KRASG12D driven tumours. Importantly, loss of wild-type *Kras* in oncogenic KRAS-driven aggressive tumours significantly alters tumour progression, metastasis while impacting tumour immune cell infiltration. This study demonstrates a suppressive role for wild-type *Kras* during colon tumour initiation and highlights the critical impact of wild-type *Kras* upon therapeutic response to MAPK and tumour progression in *Kras* mutant cancers.

**Highlights:**

- Wild-type KRAS suppresses mutant KRASG12D mediated proliferation and signalling in colorectal cancer models *in vivo*
- Concomitant loss of wild-type KRAS and activation of WNT signalling promotes mutant KRAS-driven tumour initiation.
- Wild-type KRAS promotes resistance to MAPK inhibition in KRAS mutant tumours
- Loss of wild-type KRAS inhibits metastasis of late-stage mutant KRAS colorectal cancer models.

## Introduction

It is known that oncogene allelic imbalance is a frequent event in cancer cells, known to impact key oncogene loci such as KRAS and BRAF^1^. These allelic imbalances can result from genomic gains or losses, focal amplifications at the oncogene loci or loss of heterozygosity impacting the wild-type allele. Nonetheless, the functional and therapeutic consequences of such imbalances are poorly understood. Mutations in KRAS oncogene are frequently detected in many cancers, including colorectal adenocarcinoma (CRC) and pancreatic ductal adenocarcinoma (PDAC). Hotspot mutations involving codons 12, 13, 61 and 146 of KRAS lead to constitutive activation of KRAS and hyperactivation of many downstream effector signalling pathways, including MAPK and PI3K-AKT pathways^2,3^. Biochemically, these hotspot mutations have been shown to increase the abundance of GTP-bound active KRAS, through altered GTP hydrolysis and nucleotide exchange rates based on the type of mutation^4^. Recent reports have shown that as a likely consequence of their distinct biochemical properties, each specific KRAS mutation has a unique transforming potential and tumour promoting capacity dependent on the tissue or cancer type^5–7^.

It is now well established that the frequency of KRAS mutation differs widely based on the tissue of origin (40% of CRC and 93% of pancreatic cancers)^8^. We and others have previously shown that oncogenic *Kras* cooperates with *Apc* loss in driving colorectal tumorigenesis^9–11^. The majority of studies to date have focused on gain-of-function effects of *KRAS* mutations^12^. Unfortunately, and despite the recent successes with KRASG12C isoform specific inhibitors, it is widely accepted that mutant KRAS is strongly associated with resistance to therapies, particularly those targeting upstream or downstream signalling nodes such as EGFR, MEK, PI3K and mTOR^13–15^. This said, given their position at the nexus of many key oncogenic and growth promoting signalling pathways, significant efforts are underway both to directly target mutant RAS oncoproteins, and to target signalling through the wild-type molecule^16,17^.

Recent work from several labs has highlighted the heterogeneous properties of KRAS alterations in human cancer, having successfully modelled a spectrum of KRAS mutations in relevant model systems^5–7^. In addition to this spectrum of oncogenic KRAS mutations, evaluation of specific pro- or anti-tumourigenic roles for the wild-type RAS proteins in the context of oncogenic RAS across cancer types has been controversial. It has previously been proposed that wild-type KRAS can exhibit tumour suppressive characteristics in cancer^18^. More recently, it was also shown that KRAS dimerization is required for the function of oncogenic KRAS^19^. In addition, it has become clear that copy number alterations at *KRAS* such as copy number gain at the mutant allele, loss of the heterozygosity, or broader allelic imbalance can result in enhanced fitness of cancer cell lines^1,20–22^. Indeed, it has been demonstrated that KRAS gene dosage can determine phenotypic characteristics and influence outcome of pancreatic cancer models *in vivo*^23,24^. Nonetheless, a clear understanding of the mechanistic basis of wild-type KRAS function in the processes of tumour initiation and progression of KRAS mutant tumours is yet to be defined.

Understanding the function of wild-type KRAS is essential to identify effective therapeutic strategies for oncogenic KRAS-driven cancers and potential mechanisms of resistance to KRAS targeting therapies. Here, we demonstrate a critical role for wild-type KRAS in tumour initiation and progression of mutant KRAS-driven tumours. Using genetically engineered mouse (GEM) models, we show that in the presence of an oncogenic *Kras* allele loss of the wild-type *Kras* allele augments oncogenic *Kras* signalling, leading to increased tumour initiation *in vivo*. We also demonstrate this increased tumour initiation is specific to the oncogenic KRAS mutation. Furthermore, utilizing KRAS-driven colorectal cancer models we show that loss of wild type KRAS promotes sensitivity to MEK inhibition *in vivo.* Finally, we show that deletion of wild-type *Kras* allele significantly alters the progression of advanced late-stage KRAS-driven colorectal tumours. Collectively, our studies provide new insights into KRAS biology and reveal a critical role for wild-type KRAS in the therapeutic resistance of KRAS-driven cancers.

## Results

### Allelic balance of mutant and wild-type KRAS affects homeostasis in the murine intestine

To accurately model the contribution of wild-type KRAS in CRC, we designed a genetically engineered mouse model (GEMM) that allows selective deletion of the wild-type *Kras* while expressing an oncogenic *Kras*^LSL-G12D^ allele (hereafter *Kras*^G12D^). Here, the wild-type *Kras* allele is replaced by a conditional *Kras*^flox^ allele to generate *Kras*^fl/LSL-G12D^ (hereafter referred to as *Kras*^fl/G12D^) (Figure 1a). Recombination of these alleles is targeted to the intestinal epithelium through activity of a tamoxifen-inducible Cre recombinase, expressed under the control of the *Villin* promoter (*VillinCre*^ER^) (Figure 1a). Using this model we confirmed the impact of the oncogenic *Kras*^G12D^ mutation upon intestinal epithelial homeostasis, and subsequently went on to characterise any modification of this phenotype elicited by *Kras*^fl/G12D^. Intestinal tissue was sampled from mice at 30 days post induction, with mutant KRAS found to promote enterocyte proliferation and robustly suppress Paneth cell differentiation in the small intestine as a consequence of increased MAPK signalling, consistent with previous reports^25^ (Figure 1b-e). These features are further exacerbated by deletion of the wild-type copy in *Kras*^fl/G12D^ mice, exemplified by increased proliferation in the intestinal crypt (BrdU^+^) (Figure 1d, e). Moreover, Lysozyme and periodic acid-Schiff (PAS) or Alcian blue (AB) stains indicated further suppression of the Paneth cell lineage and increased abundance of secretory goblet cells (Figure 1d, e). These *in vivo* data are suggestive of a role for wild-type KRAS in restraining the impact of oncogenic KRAS mutation on the intestinal epithelium, which in turn translates into quantitative phenotypic differences.

**Figure 1:**
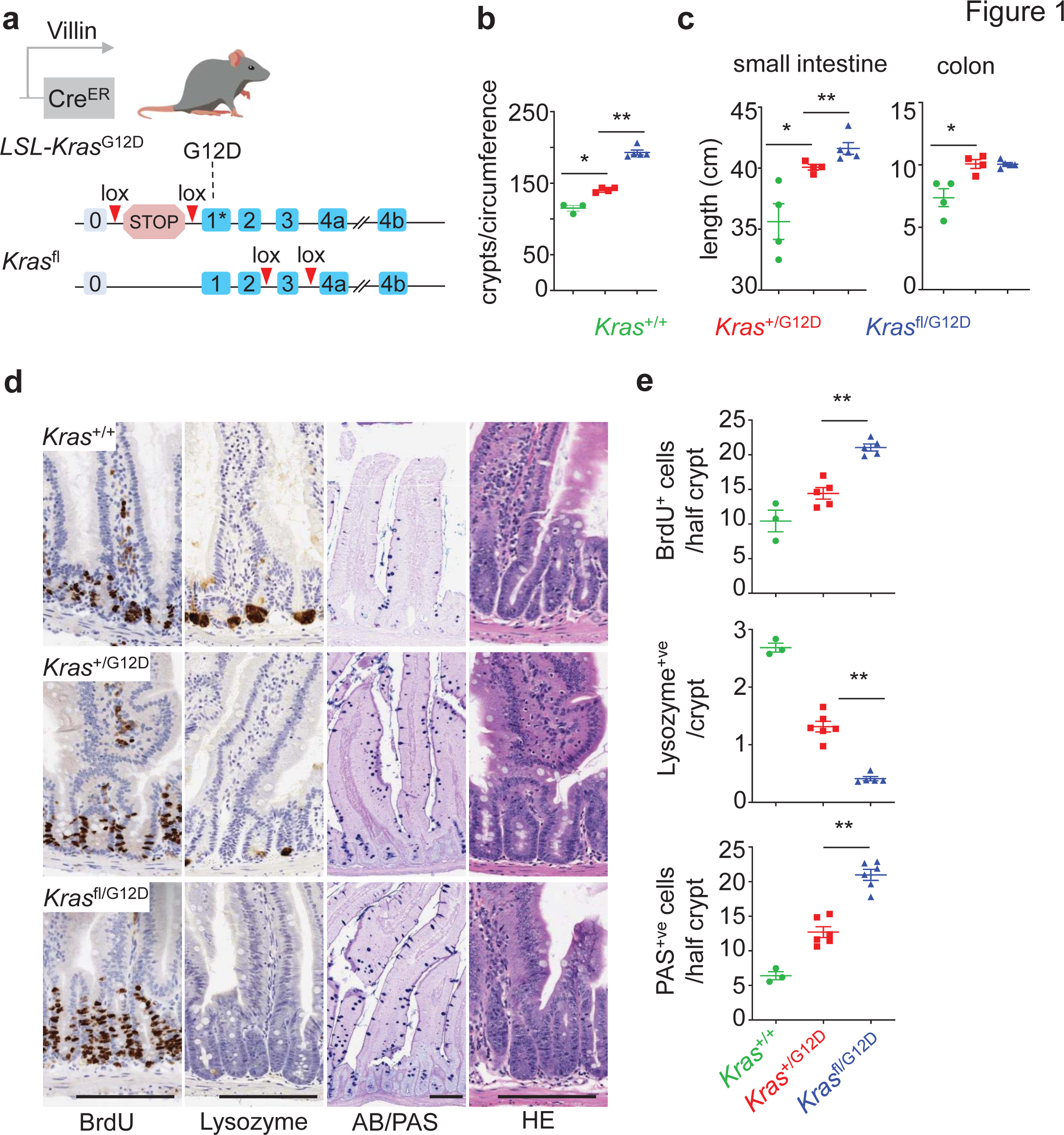
Wild-type *Kras* deletion alters *Kras*^G12D^ mutant intestinal homeostasis. a) Schematic representing the generation of *Kras^f^*^l/G12D^ mice: Villin Cre, Cre recombinase; ER, oestrogen receptor; lox, Cre-Lox recombination site; LSL, Lox-Stop-Lox cassette. b) Number of crypts per circumference from at least 25 of the small intestine of *Kras*^+/+^, *Kras*^+/G12D^ and *Kras*^fl/G12D^ mice, sampled 30 days post Cre-induction (*Kras*^+/+^, n = 3; *Kras*^+/G12D^, n = 4 and *Kras*^fl/G12D^, n = 5). Data are ± s.e.m. \**P* = 0.0286, \*\**P* = 0.0079, one-way Mann–Whitney U test. c) Length (cm) of small intestine (SI) and Colon of *Kras*^+/+^, *Kras*^+/G12D^ and *Kras*^fl/G12D^ mice, sampled 30 days post Cre-induction (*Kras*^+/+^,n = 4; *Kras*^+/G12D^, n = 4 and *Kras*^fl/G12D^, n = 5). Data are mean ± s.e.m. \**P* = 0.0143, \*\**P* = 0.0079 one-way Mann–Whitney U test. d) Representative 5-bromo-2′-deoxyuridine (BrdU), Lysozyme, Alcian blue/periodic acid– Schiff (AB/PAS) and H&E staining of *Kras*^+/+^, *Kras*^+/G12D^ and *Kras*^fl/G12D^ mouse small intestine, sampled 30 days post Cre-induction. Scale bar, 100 μm. e) Top: Number of BrdU-positive cells from at least 25 half-crypts in SI from (d). Data are mean ± s.e.m, (*Kras*^+/+^, n = 3; *Kras*^+/G12D^, n = 5; *Kras*^fl/G12D^ n = 5), \*\**P* = 0.004, one-way Mann–Whitney U test. Middle: Number of Lysozyme-positive cells from at least 25 crypts in SIof *Kras*^+/+^, *Kras*^+/G12D^ and *Kras*^fl/G12D^ in SI from (d) Data are mean ± s.e.m, (*Kras*^+/+^, n = 3; *Kras*^+/G12D^, n = 6; *Kras*^fl/G12D^ n = 5), \*\**P* = 0.0022, one-way Mann– Whitney U test. Bottom: Number of PAS-positive cells from at least 25 half-crypts in SI. *Kras*^+/+^, *Kras*^+/G12D^ and *Kras*^fl/G12D^ in SI from (d) Data are mean ± s.e.m, (*Kras*^+/+^, n = 3; *Kras*^+/G12D^, n = 6; *Kras*^fl/G12D^ n = 6). \*\**P* = 0.0011, one-way Mann–Whitney U test.

Allelic imbalance is reported to be a common feature associated with many oncogenes in addition to *KRAS*^1^. For example, BRAF allelic imbalance is reported to occur in 40% of BRAF mutant skin cancers^26^. Therefore, we tested whether *Braf* allelic imbalance could also impact intestinal homeostasis. Here, we assessed the impact of a conditional oncogenic *Braf^V^*^600^*^E^* allele, alone or in combination with a conditional *Braf* targeting allele (henceforth referred to as *Braf^fl/V^*^600^*^E^*), or when bred to homozygosity (*Braf^V^*^600^*^E/V^*^600^*^E^*), again under the control of *VillinCre^ER^*. To assess the impact of *Braf^V^*^600^*^E^* gene dosage, intestinal tissues were sampled from *Braf^+/V^*^600^*^E^*, *Braf^fl/V^*^600^*^E^*, *Braf^V^*^600^*^E/V^*^600^*^E^* or control mice at a timepoint 3 days post-induction recombination. As previously reported, we found that *Braf^V^*^600^*^E/+^* promotes proliferation in the intestinal crypt (BrdU^+^), indeed to a greater degree than *Kras^G12D/+^* at this timepoint (Extended Figure 1a, b)^27,28^. In line with the phenotypes observed in KRAS mutant intestine, we find that altering allelic balance in favour of oncogenic *Braf*, either through breeding to homozygosity (*Braf^V^*^600^*^E/V^*^600^*^E^*), or through conditional deletion of the wild-type allele (*Braf^fl/V^*^600^*^E^*) leads to significant increase in crypt cell proliferation and loss of Lysozyme-positive Paneth cells (Extended Figure 1a, b). However, BRAF activation does not significantly alter the abundance of secretory goblet cells in the intestinal epithelium (Extended Figure 1a, b). Collectively, these data show that allelic imbalance at oncogene loci akin to that seen in human cancer exacerbates specific oncogene associated and cancer related phenotypes.

### Wild-type *Kras* deletion in the presence of oncogenic KRASG12D and Wnt activation accelerates tumorigenesis via MAPK signalling

Given the observed impact upon normal intestinal homeostasis driven by altered gene dosage at oncogenic loci, we investigated whether these events might cooperate with the concomitant loss of the tumour suppressor gene *Apc*. The *Kras*^fl/G12D^ mouse line described above was bred to the well characterised mouse line bearing conditional deletion of the tumour suppressor gene *Apc* (*VillinCre*^ER^ *Apc*^fl/fl^), generating *Apc*^fl/fl^ *Kras*^fl/G12D^ or *Apc*^fl/fl^ *Kras*^+/G12D^ mouse lines as simple, tractable models of oncogene-induced hyperproliferation *in vivo.* Homozygous deletion of *Apc* in the murine intestine results in a robust phenotype, driven by hyperproliferation and altered differentiation of the intestinal crypt epithelium^29^. Moreover, we have shown that this phenotype is exacerbated by expression of oncogenic *Kras*^10,11^. Using this system, we investigated whether loss of wild-type *Kras*, in the context of an oncogenic *Kras*^G12D^ mutation and *Apc* loss, impacted epithelial proliferation *in vivo*. Indeed, this was the case with significantly enhanced proliferation, as denoted by BrdU incorporation, observed in the intestinal epithelium of *Apc*^fl/fl^ *Kras*^fl/G12D^ mice when compared to *Apc*^fl/fl^ *Kras*^+/G12D^ mice (Figure 2a, b), with the area of proliferative cells extending higher in the villus epithelium in *Apc*^fl/fl^ *Kras*^fl/G12D^ mice. This was not due to *Kras* copy number or allele changes as confirmed using droplet PCR (Extended Figure 2a, b). The ectopic proliferation/dedifferentiation of cells in the villus epithelium is a key feature of *Kras*^G12D^ mutation in the context of *Apc* deficiency, and is concomitant with an acquired ability of mutant cells to form organoid cultures *in vitro*^11^. Consistent with the observed proliferation in the villus epithelium of *Apc*^fl/fl^ *Kras*^fl/G12D^ mice, the characteristic organoid forming capacity was enhanced when compared to *Apc*^fl/fl^ *Kras*^+/G12D^ mice (Extended Figure 2c, d). These data indicate that loss of the wild-type copy of *Kras* in the context of concomitant oncogenic *Kras*^G12D^ mutation and *Apc* depletion can enhance a number of *Kras*^G12D^ associated phenotypes.

**Figure 2:**
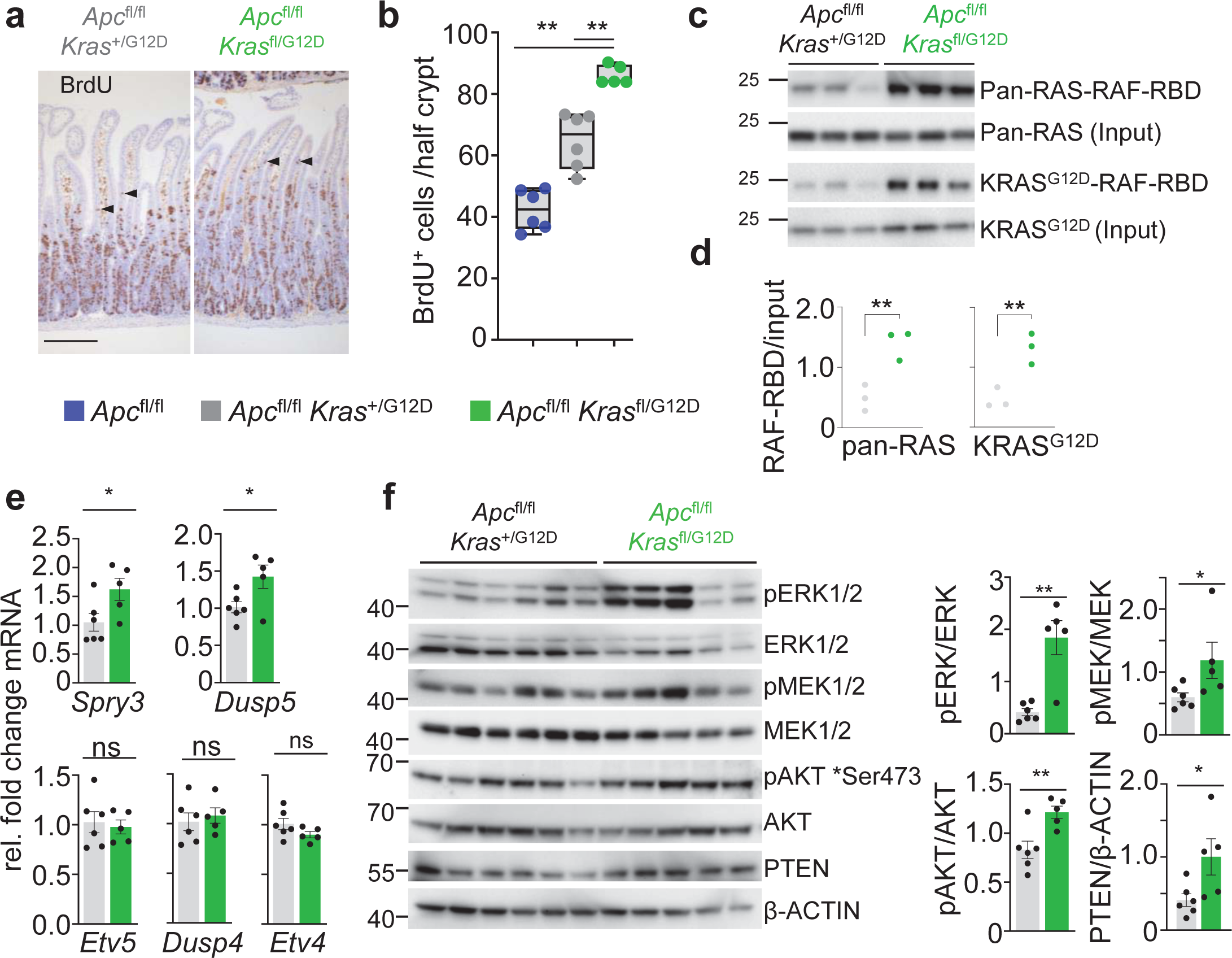
Wild-type *Kras* deletion increases proliferation, active KRAS and MAPK signaling. a) Representative H&E and BrdU IHC of *Apc*^fl/fl^ *Kras*^fl/G12D^ mice sampled 3 days post Cre-induction. Dashed boxes highlight selected area shown in higher magnification. Arrow heads indicate de-differentiating cells in the villi. Scale bar, 100 μm. b) Boxplots showing BrdU-positive cells from at least 25 half-crypts in SIin *Apc*^fl/fl^*, Apc*^fl/fl^ *Kras*^+/G12D^ and *Apc*^fl/fl^ *Kras*^fl/G12D^ of mice sampled 3 days post Cre-induction. Boxes depict interquartile range, central line indicates median and whiskers indicate minimum/maximum values *(Apc*^fl/fl^,n *=* 6*; Apc*^fl/fl^ *Kras*^+/G12D^, n = 6; and *Apc*^fl/fl^ *Kras*^fl/G12D^, n = 5 mice). \*\**P* = 0.0022, \*\**P* = 0.0043, one-way Mann–Whitney U test. c) RAF-Ras Binding Domain (RBD) agarose affinity purification assay of three biologically independent samples per condition from *Apc*^fl/fl^ *Kras*^+/G12D^ *and Apc*^fl/fl^ *Kras*^fl/G12D^ intestinal organoids. Pulldown of RAS-GTP with RAF-RBD agarose beads. Top: Precipitates were immunoblotted using a pan-RAS antibody and input pan-RAS served as loading control. RAS-GTP activation levels were quantified and normalised to pan-Ras loading control. Bottom: Precipitates were immunoblotted using a KRAS^G12D^ antibody and input KRAS^G12D^ served as loading control. KRAS^G12D^-RAF-RBD levels were quantified and normalised to KRAS^G12D^ loading control. 3 biological replicates per group, each lane represents organoids generated from individual mice from the genotype indicated. d) Quantification of RAF-RBD assay (left) and KRASG12D immunoblot (right) from c), \*\**P* = 0.0045 (pan-RAS), **P = 0.0044 (KRAS^G12D^), one-way Mann–Whitney U test. e) qRT-PCR analysis of *Etv5, Dusp4, Etv4* and *Spry3* in *Apc*^fl/fl^ *Kras*^+/G12D^ *and Apc*^fl/fl^ *Kras*^fl/G12D^ intestinal organoids. Transcript levels were normalised to *Gapdh* (n = 6 independent organoids per group). \**P* < 0.05 (Etv5, *Dusp4*), one-way Mann–Whitney U test. f) Left: Immunoblots showing PTEN, pAKT (Ser473), AKT, pERK1/2, ERK1/2, pMEK1/2, MEK1/2 and ß-actin in *Apc*^fl/fl^ *Kras*^+/G12D^ *and Apc*^fl/fl^ *Kras*^fl/G12D^ intestinal organoids. 6 *Apc*^fl/fl^ *Kras*^+/G12D^ *and* 5 *Apc*^fl/fl^ *Kras*^fl/G12D^ biological replicates per group, each lane represents organoids generated from individual mice from genotype indicated. Right: Quantification of immunoblots, phosphorylated proteins were normalised to total protein levels. \*\**P* = 0.0043 (pERK/ERK), \**P* = 0.0152 (pMEK/MEK), \*\**P* = 0.0087 (pAKT/AKT), \**P* = 0.015 (PTEN/ ß-actin), one-way Mann–Whitney U test.

We hypothesized that loss of the wild-type *Kras* allele may alter baseline gene dosage of the oncogenic mutant allele, and as a result, lead to enhanced KRAS activation and increased signalling flux through downstream effector pathways and any associated feedback loops. This hypothesis was tested through comparison of organoid cultures derived from *Apc*^fl/fl^ *Kras*^fl/G12D^ and *Apc*^fl/fl^ *Kras*^+/G12D^ intestinal tissues. Initially, we quantified the relative proportion of GTP-bound, active RAS proteins. In these pull-down assays, *Apc*^fl/fl^ *Kras*^fl/G12D^ organoids were characterised by enhanced binding of RAS to the RAS-binding domain (RBD) of BRAF, when compared to *Apc*^fl/fl^ *Kras*^+/G12D^, suggestive of a larger pool of active RAS (Figure 2c, d). It is well known that increased RAS activity translates into increased activation of downstream effector pathways such as the MAPK cascade^30^. To determine whether MAPK activity was increased in *Apc*^fl/fl^ *Kras*^fl/G12D^ and *Apc*^fl/fl^ *Kras*^+/G12D^ organoids, we quantified expression of known ERK-regulated transcripts, finding these to be enriched in *Apc*^fl/fl^ *Kras*^fl/G12D^ organoids (Figure 2e). To evaluate MAPK activity in more depth, we assessed downstream effector activity in these lines through quantitative immunoblotting. We detected a clear increase in phosphorylation of ERK1/2 and MEK1/2 in *Apc*^fl/fl^ *Kras*^fl/G12D^ organoids, while phosphorylation of AKT and expression of PTEN were unaffected (Figure 2f). These data indicate that loss of the wild-type copy of *Kras* enhances the relative activity of the oncogenic mutant allele and drives downstream effector pathway signalling.

Given the robust impact of wild-type *Kras* deletion in the acute setting above, we next addressed the role of wild-type KRAS on oncogenic KRASG12D-driven intestinal tumorigenesis. To this end, we generated *VillinCre*^ER^ *Apc*^fl/+^ *Kras*^+/G12D^ (henceforth *AKras*^+/G12D^) and *VillinCre*^ER^ *Apc*^fl/+^ *Kras*^fl/G12D^ (henceforth *AKras*^fl/G12D^). In this setting, intestinal tumour development occurs following sporadic loss of the second copy of *Apc* in individual intestinal crypts. Targeted mutation in the intestinal epithelium was induced through intraperitoneal administration of tamoxifen, with deletion of wild-type *Kras* in the context of oncogenic *Kras*^G12D^ (*AKras*^fl/G12D^) resulting in a significant acceleration of tumorigenesis, and a consequent reduction of median survival based upon an endpoint defined by clinical signs associated with tumour burden. (Figure 3a). The reduced time to onset of signs associated with intestinal tumorigenesis in *AKras*^fl/G12D^ mice was coincident with a striking tumour initiation phenotype, with development of numerous small lesions observed principally in the small intestine (Figure 3b). The initiating lesions observed in *AKras*^fl/G12D^ mice were histopathologically comparable to those observed in *AKras*^+/G12D^ mice (Figure 3c), and as expected, were positive for nuclear β-catenin (Figure 3d). Immunohistochemical analysis demonstrated nuclear accumulation of phosphorylated ERK1/2 (Figure 3d), increased expression of c-MYC and increased abundance of γH2AX in *AKras*^fl/G12D^ tumours when compared to AKras+/G12D tumours, suggestive of MAPK pathway activation, increased cellular proliferation and activation of DNA damage response pathways (Figure 3e, f). Indeed, using BrdU incorporation as a marker for cellular proliferation, we demonstrated that the tumour epithelium of *AKras*^fl/G12D^ mice is markedly more proliferative than that of *AKras*^+/G12D^ mice (Figure 3g). Together, these results show that loss of the wild-type copy of *Kras* increases activity and signalling of the oncogenic mutant allele driving tumour initiation.

**Figure 3:**
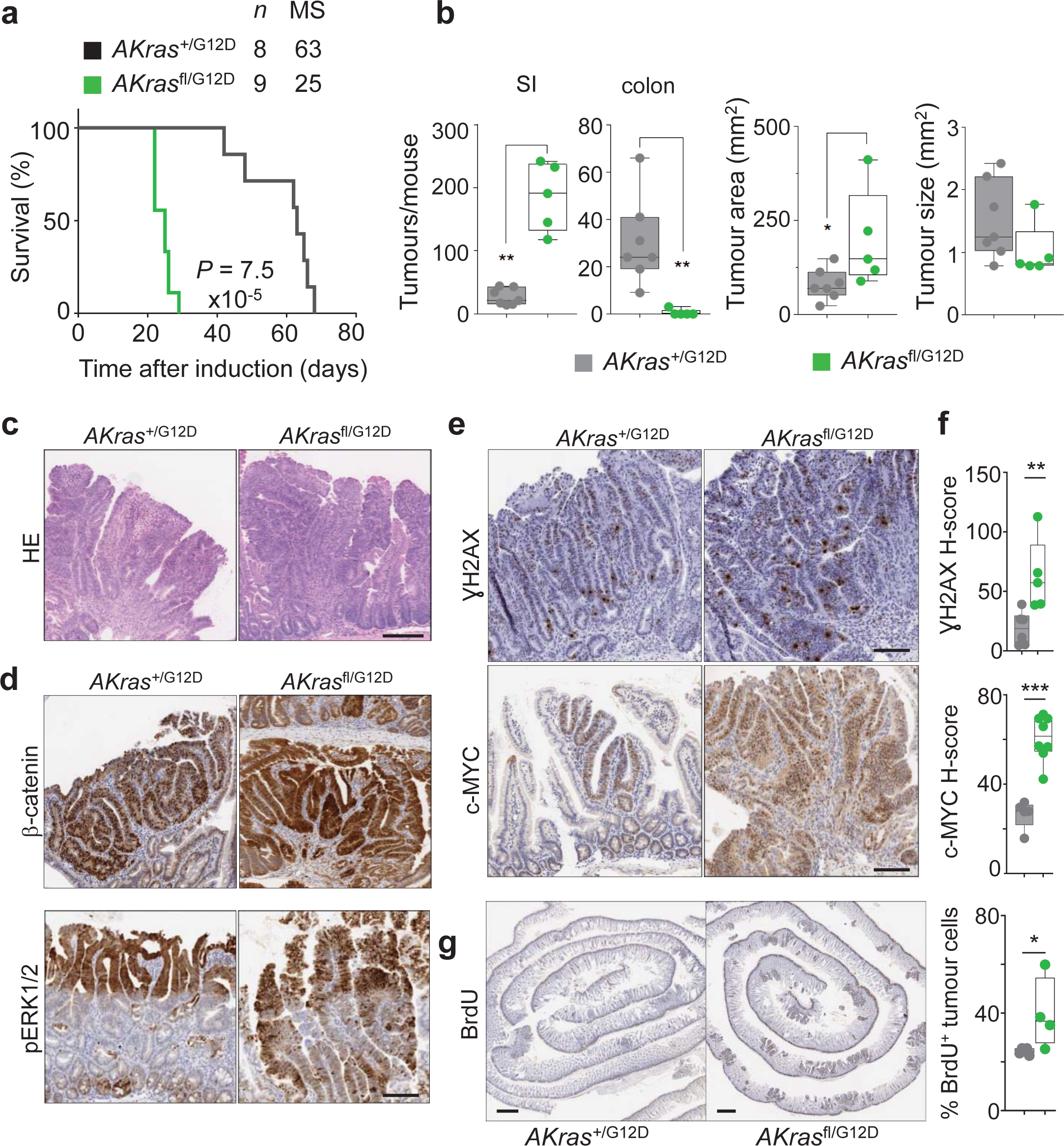
Loss of wild-type *Kras* increases mutant KRAS-driven tumourigenesis together with *Apc* loss. a) Kaplan-Meier survival curve of *VillinCre^ER^ Apc^+^*^/fl^ *Kras*^+/G12D^ (A*Kras*^+/G12D^) and *VillinCre^ER^ Apc*^+/fl^ *Kras*^fl/G12D^ (A*Kras*^fl/G12D^) mice aged until clinical endpoint (*Apc^+^*^/fl^ *Kras*^+/G12D^,n = 8, MS = 63; *Apc*^+/fl^ *Kras*^fl/G12D^, n = 9 mice, MS = 25), \*\*\*\**P* = 7.5 x10^-5^, log-rank (Mantel-Cox) test. b) Left: Boxplots showing total number of tumours from *APC*^+/fl^ *Kras*^+/G12D^ and *Apc*^+/fl^ *Kras*^fl/G12D^ mice aged until clinical endpoint in SI and Colon. Right: Boxplots showing tumour area (mm^2^) and tumours size (mm^2^) in *Apc*^+/fl^ *Kras*^+/G12D^ and *Apc*^+/fl^ *Kras*^fl/G12D^ mice aged until clinical endpoint. Boxes depict interquartile range, central line indicates median and whiskers indicate minimum/maximum values (*Apc*^+/fl^ *Kras*^+/G12D^, n = 7; *Apc*^+/fl^ *Kras*^fl/G12D^ n= 5 mice). \**P* < 0.05, \*\**P* < 0.01 one-way Mann–Whitney U test. c) Representative H&E images of *Apc*^+/fl^ *Kras*^fl/G12D^ tumour. Dashed box highlights selected area shown in high magnification. d) Representative images of nuclear β-catenin and pERK1/2 staining in *Apc*^+/fl^ *Kras*^+/G12D^ and *Apc*^+/fl^ *Kras*^fl/G12D^ tumours at clinical endpoint. Representative of 5 biological replicates. Scale bars, 100 μm. e) Representative images of γH2AX and c-MYC staining in *Apc*^+/fl^ *Kras*^+/G12D^ and *Apc*^+/fl^ *Kras*^fl/G12D^ tumours at clinical endpoint. Representative of 5 biological replicates. Scale bars, 100 μm. f) H-score of γH2AX and c-MYC IHC staining of e). Boxes depict interquartile range, central line indicates median and whiskers indicate minimum/maximum values, (*Apc*^+/fl^ *Kras*^+/G12D^, n = 5; *Apc*^+/fl^ *Kras*^fl/G12D^ n = 5 mice). \*\**P* = 0.0043, \*\*\**P* = 0.0008, one-way Mann–Whitney U test. g) Representative BrdU staining of *Apc*^+/fl^ *Kras*^+/G12D^ and *Apc*^+/fl^ *Kras*^fl/G12D^ small intestinal tumours at clinical endpoint. Scale bars, 100 μm. Right: quantification of BrdU positivity in tumour cells. Boxes depict interquartile range, central line indicates median and whiskers indicate minimum/maximum values. (*Apc*^+/fl^ *Kras*^+/G12D^, n = 4; *Apc*^+/fl^ *Kras*^fl/G12D^ n = 4 mice). \**P* < 0.05, one-way Mann–Whitney U test.

### Loss of wild-type KRAS restores sensitivity to MEK inhibition in colorectal tumours *in vivo*

We have demonstrated that altered allelic balance of oncogenic *Kras*^G12D^ has a substantial impact upon tumour initiation in models of intestinal disease. Given that *KRAS* mutation is clinically associated with resistance to targeted therapies, we next investigated whether therapeutic efficacy is positively or negatively influenced by allelic imbalance at the *Kras* locus. We have previously shown that the observed intestinal crypt epithelium hyperproliferation characteristic of the *Apc*^fl/fl^ *Kras*^+/G12D^ model is resistant to MEK inhibition^11,31^, and that this proliferation is enhanced in *Apc*^fl/fl^ *Kras*^fl/G12D^ mice concomitant with increased KRAS and MAPK activity (Figure 2c,f). We reasoned that this increased MAPK activation might result in an acquired sensitivity to inhibition of MEK1/2 with a clinically relevant targeted therapeutic agent (AZD6244/selumetinib). As previously, we found that treatment of *Apc*^fl/fl^ *Kras*^+/G12D^ mice with AZD6244 had no impact upon intestinal hyperproliferation. Importantly, and in contrast, treatment with AZD6244 not only significantly decreased proliferation in the intestinal crypt epithelium of *Apc*^fl/fl^ *Kras*^fl/G12D^ mice but suppressed proliferation to a level below that of vehicle treated *Apc*^fl/fl^ *Kras*^+/G12D^ mice (Figure 4a). We next tested whether MEK1/2 inhibition had a similar suppressive effect upon the process of intestinal tumourigenesis. To do this, we treated *AKras*^fl/G12D^ or *AKras*^+/G12D^ mice with AZD6244 (25mgkg^-1^, BID) from 1-day post-induction of genetic recombination (Figure 4b). We found that inhibition of MEK1/2 had a modest impact on the survival of *AKras*^+/G12D^ mice, based upon endpoint defined by onset of clinical signs associated with tumour burden, but significantly extended survival in *AKras*^fl/G12D^ mice (median survival extended from 26 days to 115 days) (Figure 4c). This extension in survival was accompanied by a dramatic reduction in the number of small intestinal tumours (Figure 4d) alongside increase in colonic tumour number, albeit at greatly increased time post induction. A significant reduction in tumour cell proliferation (BrdU incorporation) was also observed in *AKras*^fl/G12D^ mice compared to *AKras*^+/G12D^ derived tumours (Figure 4e). These data suggest that wild-type *Kras* acts to suppress the penetrance of mutant oncogenic *Kras*^G12D^, thus dampening MAPK signalling and contributing to therapeutic resistance in *Kras* mutant tumours, a key clinical problem.

**Figure 4:**
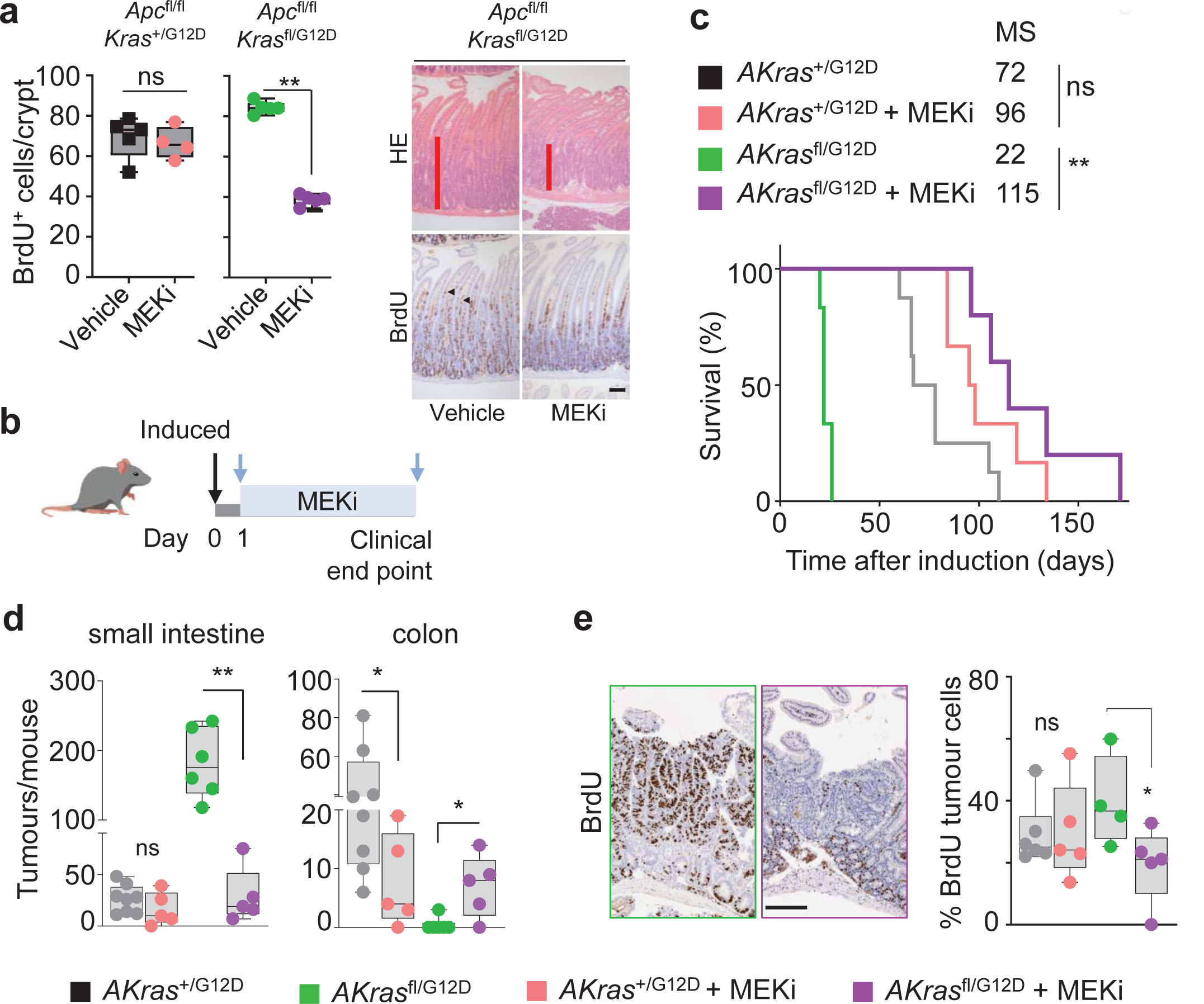
Lack of wild-type *Kras* increases sensitivity to MEK inhibition in of *Kras*^G12D^ colorectal tumours *in vivo*. a) Left, quantification of BrdU positive cells per half crypt in *Apc*^fl/fl^ *Kras*^+/G12D^ and *Apc*^fl/fl^ *Kras*^fl/G12D^ mice 3 days post-induction treated with Vehicle or MEKi (AZD6244) as indicated. Boxes depict interquartile range, central line indicates median and whiskers indicate minimum/maximum values (*Apc*^fl/fl^ *Kras*^+/G12D^ Vehicle, n = 5; MEKi, n = 4; and *Apc*^fl/fl^ *Kras*^fl/G12D^ Vehicle, n= 5; MEKi, n = 5). \*\**P* = 0.004, one-way Mann–Whitney U test. Right, representative H&E and BrdU images of *Apc*^fl/fl^ *Kras*^fl/G12D^ 3 days post-induction treated with Vehicle or MEKi (AZD6244) as indicated. Arrow heads show BrdU^+ve^ hyper proliferative cells. Scale bar, 100 μm b) Schematic presenting experimental approach. *Apc*^+/fl^ *Kras*^+/G12D^ and *Apc*^+/fl^ *Kras*^fl/G12D^ mice treated with MEKi one day post-induction and treated to clinical endpoint. c) Kaplan-Meier survival curve of *Apc*^+/fl^ *Kras*^+/G12D^ and *Apc* ^+/fl^ *Kras*^fl/G12D^ mice treated as shown in b) aged until clinical endpoint (*Apc* ^+/fl^ *Kras*^+/G12D^, n = 8, MS = 72; *Apc* ^+/fl^ *Kras*^+/G12D^ MEKi, n = 6, MS = 96; *Apc*^+/fl^ *Kras*^fl/G12D^,n = 6, MS = 26; *Apc* ^+/fl^ *Kras*^fl/G12D^ MEKi, n = 5, MS = 115). ** *P* = 0.0014, ns = not significant, log-rank (Mantel-Cox) test. d) Boxplots showing total number of tumours from *Apc*^+/fl^ *Kras*^+/G12D^ and *Apc*^+/fl^ *Kras*^fl/G12D^ mice untreated or treated with MEKi as indicated in c and aged until clinical endpoint in SI and Colon. Boxes depict interquartile range, central line indicates median and whiskers indicate minimum/maximum values *(Apc*^+/fl^ *Kras*^+/G12D^, n = 8; *Apc*^+/fl^ *Kras*^+/G12D^ MEKi, n = 5; *Apc*^+/fl^ *Kras*^fl/G12D^,n = 6; *Apc*^+/fl^ *Kras*^fl/G12D^ MEKi, n = 5). \*\**P* = 0.0022 (*Apc*^+/fl^ *Kras*^fl/G12D^ SI), \**P* = 0.0281, * *P* = 0.0152 (colon), one-way Mann–Whitney U test. e) Left: Representative BrdU images of *Apc*^+/fl^ *Kras*^+/G12D^ and *Apc*^+/fl^ *Kras*^fl/G12D^ tumours from mice treated with MEKi from day 1 post-induction until clinical endpoint. Right: boxplot showing percentage of BrdU positive tumour cells in *Apc* ^+/fl^ *Kras*^+/G12D^ and *Apc* ^+/fl^ *Kras*^fl/G12D^ treated with MEKi. Boxes depict interquartile range, central line indicates median and whiskers indicate minimum/maximum values, (*Apc* ^+/fl^ *Kras*^+/G12D^ untreated, n = 6; *Apc*^+/fl^ *Kras*^+/G12D^ MEKi, n = 4; *Apc* ^+/fl^ *Kras*^fl/G12D^ untreated, n = 4; *Apc*^+/fl^ *Kras*^fl/G12D^ MEKi, n = 5), \**P* = 0.0159, one-way Mann–Whitney U test.

### Loss of wild-type KRAS potentiates oncogenic *Kras*^G12D^ driven tumorigenesis in the absence of exogenous WNT mutations *in vivo*

Within the intestinal epithelium KRAS mutation alone is not sufficient to drive tumourigenesis. Indeed, in human, KRAS mutant clones are known to commonly arise with age, in the intestinal epithelium in morphologically normal crypts. This is recapitulated by murine models where expression of an oncogenic *Kras*^G12D^ mutant allele in isolation throughout the intestinal epithelium (*VillinCre*^ER^ *Kras*^+/G12D^) results in lowly penetrant intestinal lesion development, which translates to an overall survival of greater than 12 months. Given the impact of combined deletion of wild-type *Kras* and expression of oncogenic *Kras*^G12D^ on both intestinal homeostasis, and tumour initiation in the context of heterozygous loss of *Apc*, we next asked whether these alterations in *Kras* allelic balance could drive tumour initiation in isolation. Here, we induced genetic recombination in *VillinCre*^ER^ *Kras*^fl/G12D^ or *VillinCre*^ER^ *Kras*^+/G12D^ mice, and assessed survival based upon sampling due to onset clinical signs associated with significant intestinal tumour burden. As expected, *VillinCre*^ER^ *Kras^+/G12D^* alone resulted in inefficient tumour initiation, with median survival for this group at around 16 months. Strikingly, mice in the *VillinCre*^ER^ *Kras*^fl/G12D^ group developed tumours much more rapidly than the control group, exhibiting a greatly reduced median survival of around 230 days/9 months (Figure 5a). The tumours from both *Kras*^fl/G12D^ and *Kras*^+/G12D^ mice were predominantly found in the small intestine, and were characteristically large adenomas. However, tumour number was significantly increased in the small intestine of *Kras*^fl/G12D^ mice (Figure 5b, c). This demonstrates that even when the only exogenously introduced driver mutation is *Kras*^G12D^, loss of wild-type *Kras* significantly promotes intestinal tumourigenesis, which in turn translates into significantly shortened survival.

**Figure 5:**
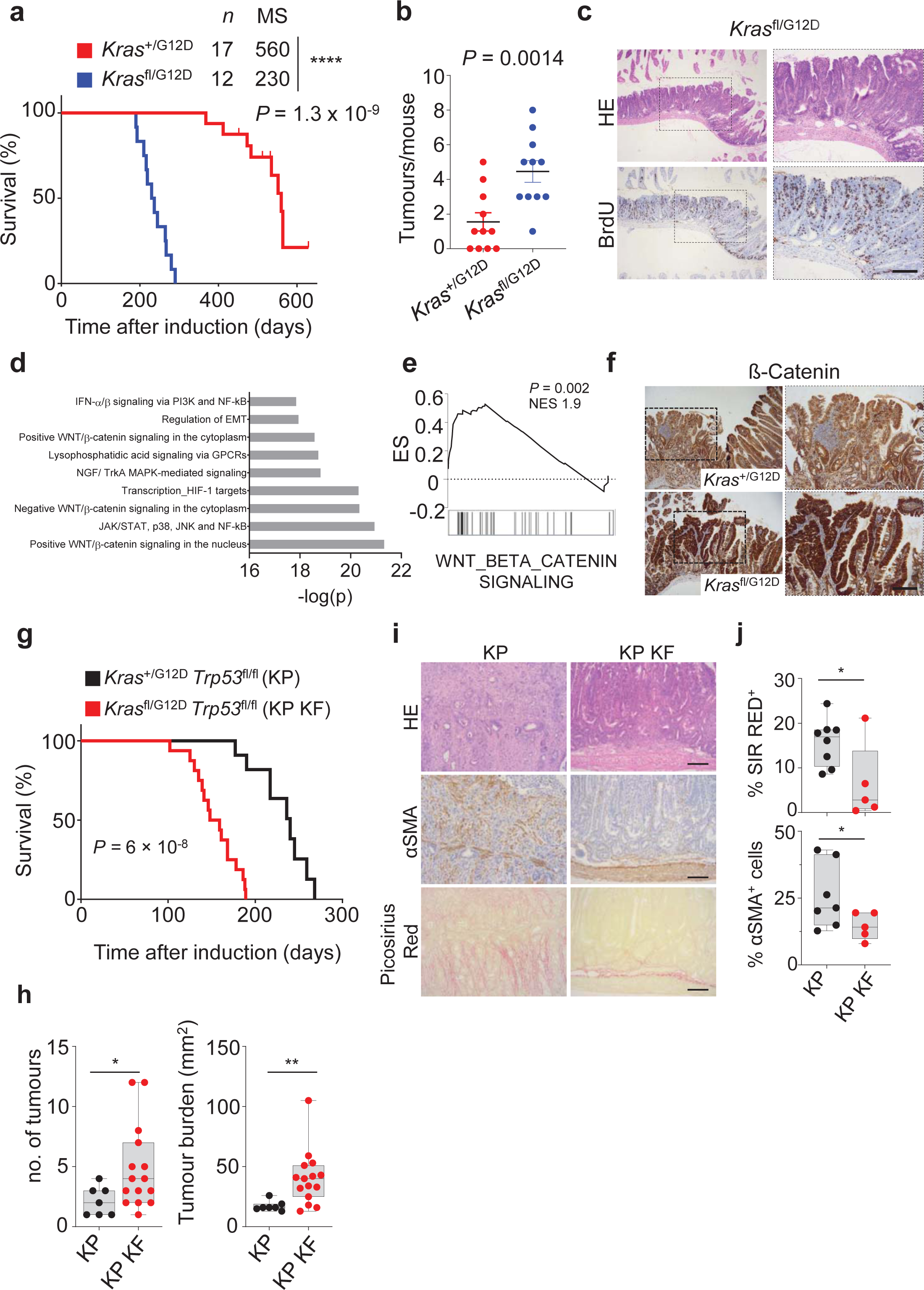
Wild-type *Kras* deletion increases tumour intiation and alters progression of *Kras*^G12D^ mutant colorectal tumours following loss of p53. a) Kaplan-Meier survival curve of *Villin*^CreER^ *Kras*^+/G12D^ and *Kras*^fl/G12D^ mice aged until clinical endpoint (*Kras*^+/G12D^, n = 17, MS = 560; *Kras*^fl/G12D^, n=12, MS = 230). \*\*\*\**P* = 1.3 x 10^-9^, log-rank (Mantel-Cox) test. b) Graph showing total number of tumours from *Kras*^+/G12D^ and *Kras*^fl/G12D^ mice. Data are mean ± s.e.m. (*Kras*^+/G12D^, n = 11; *Kras*^fl/G12D^, n = 11). \*\**P* = 0.0014, one-way Mann– Whitney U test. c) Representative H&E and BrdU IHC images of intestine from *Kras*^fl/G12D^ mice. Dashed boxes highlight selected areas shown in high magnification. Scale bar, 200 μm. d) Metacore Network analysis of differentially expressed genes from *Kras*^fl/G12D^ tumours. e) Geneset enrichment plot for Wnt beta-catenin signalling signature from the ‘Hallmark’ gene set collection of tumours derived from *Kras*^fl/G12D^ mice. \*\**P* = 0.002, NES, normalised enrichment score. f) Representative IHC for β-catenin intestine staining in *Kras*^+/G12D^ and *Kras*^fl/G12D^ mice aged until clinical endpoint. Dashed boxes highlight selected areas shown in high magnification. Scale bars, 200 μm. g) Kaplan-Meier survival curve of *Kras*^+/G12D^ *Trp53*^fl/fl^ (KP) and *Kras*^fl/G12D^ *Trp53*^fl/fl^ (KP KF) mice aged until clinical endpoint (KP, n = 19; KP KF, n = 16) \*\*\*\**P* = 6×10^-8^, log-rank (Mantel-Cox) test. h) Boxplots showing total number of tumours (left) and tumour burden (mm^2^) right in *Kras*^+/G12D^ *Trp53*^fl/fl^ (KP) and *Kras*^fl/G12D^ *Trp53*^fl/fl^ (KP KF) mice aged until clinical endpoint. Boxes depict interquartile range, central line indicates median and whiskers indicate minimum/maximum values (KP, n = 7, MS = 240; KP KF, n = 15, MS = 153). \**P* =0.02, ** *P* = 0.0017, one-way Mann-Whitney U test. i) Representative H&E, αSMA (alpha-smooth muscle actin) and Sirius Red staining of *Kras*^+/G12D^ *Trp53*^fl/fl^ (KP) *a*nd *Kras*^fl/G12D^ *Trp53*^fl/fl^ (KP KF) tumours. Representative of KP n = 8; KP KF n = 5. Scale bar, 100 μm. j) Boxplots showing Sirius Red positivity (%) and αSMA positive cells (%) of *Kras*^+/G12D^ *Trp53*^fl/fl^ (KP) and *Kras*^fl/G12D^ *Trp53*^fl/fl^ (KP KF) tumours. Boxes depict interquartile range, central line indicates median and whiskers indicate minimum/maximum values (KP n = 7; KP KF, n = 5). *\*P* = 0.0326 (Sirius Red)*, *P =* 0.024 (αSMA), one-way Mann–Whitney U test.

To better understand the mechanistic impact of deletion of wild-type *Kras* in the context of oncogenic *Kras*^G12D^, we transcriptionally profiled tumours arising in *VillinCre*^ER^ *Kras*^fl/G12D^ mice and compared to adjacent non-transformed tissue. Geneset enrichment analysis (GSEA), identified key oncogenic programmes associated with KRAS signalling, MEK and AKT were enriched in *VillinCre*^ER^ *Kras*^fl/G12D^ tumours (Extended Figure 3a). Importantly, GSEA and Metacore analysis of differentially expressed genes showed an overrepresentation of WNT and β-catenin signalling pathways (Figure 5d, e). Given that aberrant activation of the Wnt/β-catenin pathway, and its downstream transcriptional networks, is a common initiating event in CRC, and results from stabilization and nuclear accumulation of the transcriptional co-activator β-catenin, we interrogated tumours arising in *VillinCre*^ER^ *Kras*^fl/G12D^ and *VillinCre*^ER^ *Kras*^+/G12D^ mice for nuclear accumulation of β-catenin. Consistent with the transcriptional enrichment of Wnt signalling pathways (Figure 5e), tumours arising in *Kras*^fl/G12D^ intestines showed positivity for nuclear β-catenin while *Kras*^+/G12D^ intestines more predominantly exhibited membranous β-catenin (Figure 5f).

### Loss of wild-type KRAS alters tumour progression and metastasis of aggressive mutant KRAS-driven colorectal tumours

Thus far, we have only investigated the role of wild type KRAS in mouse models of adenoma. Recent reports have shown that changes in *KRAS* gene dosage alter the clonal evolution of tumours and change the metastatic incidence in KRAS mutant cancers^23^. Given the dramatic role we have observed in tumour initiation, we next wished to see how important wild-type KRAS would be in tumour progression. We have previously demonstrated that deletion or mutation of the tumour suppressor gene *Trp53* alongside activating mutation of *Kras* can give rise to aggressive, late-stage adenocarcinoma development in the intestine^32^. Indeed, the human paralogue, TP53 is inactivated in more than half of human colorectal cancers (TCGA). These models therefore represent an ideal system for interrogating the impact of wild-type *Kras* deletion in a more complex, aggressive, clinically relevant oncogenic KRAS^G12D^ driven disease.

To achieve this, we interbred the *VillinCre*^ER^ *Kras*^fl/G12D^ mouse line with a *Trp53* conditional knockout allele to generate *VillinCre*^ER^ *Kras*^fl/G12D^ *Trp53*^fl/fl^ (henceforth referred to as KP KF) and *VillinCre*^ER^ *Kras*^+/G12D^ *Trp53*^fl/fl^ (henceforth referred to as KP) mice. While KP mice developed on average one intestinal tumour and exhibited a median overall survival of 240 days, with very lowly penetrant metastasis (Figure 5g, h), KP KF mice developed on average four intestinal tumours and exhibited a median overall survival of 150 days (Figure 5g, h). Histological analysis of tumours arising in KP KF mice suggested a morphology akin to human tubulovillous adenoma, in contrast to the tumours which arose in control KP mice, which exhibit a serrated morphology. Moreover, evidence of local invasion or poor differentiation, features typically associated with advanced disease, were apparent in KP tumours but absent from KP KF tumours (Figure 5i). In addition, there were clear differences in the stromal microenvironment of the KP KF tumours, such as a lack of infiltrating alpha smooth muscle actin (αSMA)-positive stromal cells and low levels of stromal collagen deposition (as indicated by picrosirius red), again, contrasted by KP tumours (Figure 5i, j). These data suggest that while loss of the wild-type copy of *Kras* in an aggressive *Kras* mutant driven model of intestinal cancer can drive accelerated tumour initiation, it does not endow tumours with increased invasion or aggression, indeed these features appear suppressed.

Considering this, we decided to further investigate the role of wild-type *Kras* in aggressive, late-stage KRAS-mutant CRC using our recently described metastatic KPN model (*VillinCre*^ER^ *Kras*^+/G12D^ *Trp53*^fl/fl^ *Rosa26*^N1icd/+^). These KPN mice develop intestinal adenocarcinoma and exhibit highly penetrant metastasis, predominantly to the liver^32^. We interbred KPN mice with KP KF mice to generate (*VillinCre*^ER^ *Kras*^fl/G12D^ *Trp53*^fl/fl^ *Rosa26*^N1icd/+^; henceforth referred to as KPN KF). Strikingly, tumorigenesis was significantly accelerated in KPN KF mice with deletion of wild-type *Kras*, reflected by a significant reduction in overall survival based upon endpoint defined by clinical signs associated with tumour burden (Figure 6a). Using droplet PCR we confirmed this acceleration in tumorigenesis of KPN KF mice was not due mutant *Kras* induced changes in *Kras* copy number (Extended Figure 6a, b). Comparative histological analysis of tumours from KPN and KPN KF mice showed that tumours arising from KPN KF showed lack of local invasion with remarkable lack of metastasis incidence (Figure 6b, c). To gain a better mechanistic insight into the impact of *Kras* deletion in this setting, we first performed transcriptional analysis of tumours derived from KPN KF and KPN mice. Interestingly, KPN KF tumours showed a significant enrichment of Wnt signalling pathway (Figure 6d) and expression of the surrogate Wnt marker Notum, compared to KPN tumours (Figure 6e, f). It is notable that relatively low levels of Wnt activation is a key feature of the highly metastatic KPN tumours, while comparable high-Wnt APC-deficient, APN tumours are non-metastatic^32^. Consistent with the high Wnt activation signature in KPN KF tumours, comparative analysis of several published Wnt transcriptional signatures in tumours from APN, KPN and KPN KF mice clearly showed higher Wnt activation in KPN KF tumours (Extended Figure 4c). Collectively, these data show that *Kras* mutant tumours lacking wild-type *Kras* in a KPN setting activate more robust levels of Wnt signalling during tumorigenesis. This said, it is noteworthy that primary tumours arising in the intestine of KPN KF mice are adenomatous, while those arising in KPN mice are of a more aggressive adenocarcinoma/carcinomatous morphology. While the underlying genetics will undoubtedly play a key role in defining the molecular makeup of the disease, the reduced histopathological grade of tumours found in KPN KF may directly influence immune and inflammatory infiltrate.

**Figure 6:**
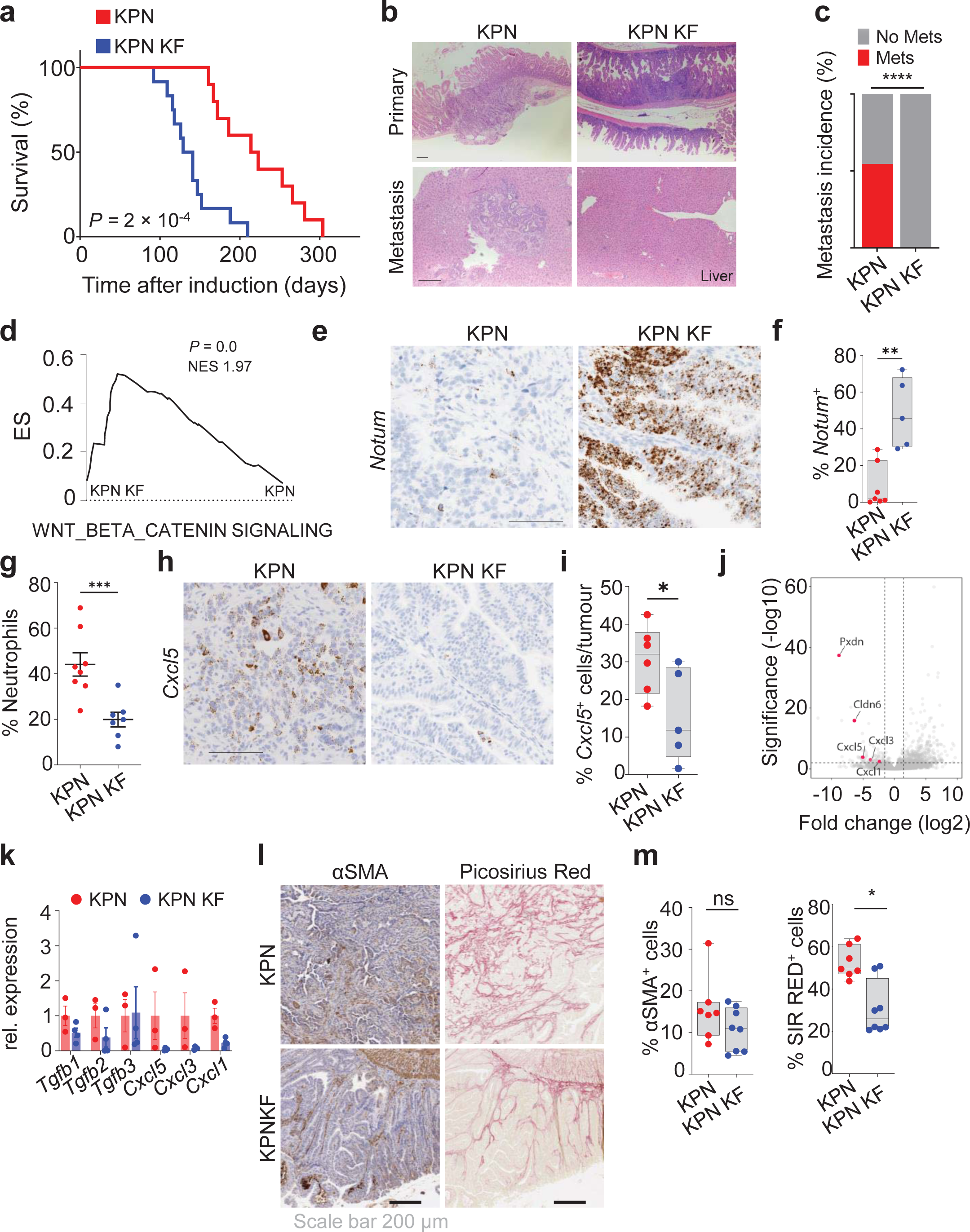
Wild-type *Kras* deletion alters progression and metastasis of KPN tumours. a) Kaplan-Meier survival curve of *Kras*^+/G12D^ *Trp53*^fl/fl^ *Rosa26*^N1icd/+^ (KPN) and *Kras*^fl/G12D^ *Trp53*^fl/fl^ *Rosa26*^N1icd/+^ (KPN KF) mice aged until clinical endpoint (KPN, n = 10, MS = 218; KPN KF, n = 12, MS = 135) \*\*\**P* = 2×10^-4^, log-rank (Mantel-Cox) test. b) Representative H&E images of primary tumour and metastasis (liver) from KPN and KPN KF mice aged to clinical endpoint. Scale bar, 200 μm. c) Incidence of metastasis (%) in *Kras*^+/G12D^ *Trp53*^fl/fl^ *Rosa26*^N1icd/+^ (KPN) and *Kras*^fl/G12D^ *Trp53*^fl/fl^ *Rosa26*^N1icd/+^ (KPN KF) mice aged until clinical endpoint (KPN, n = 10; KPN KF, n= 12). d) GSEA of hallmark WNT-β-Catenin signalling in KPN KF tumours compared to KPN. NES = normalised enrichment score. e) Representative *Notum in situ* hybridisation (ISH) in KPN and KPN KF tumours. Representative of five mice per genotype. Scale bar, 100 μm. f) Boxplots showing *Notum* (ISH) positive cells per tumour (%) of KPN and KPN KF tumours. Boxes depict interquartile range, central line indicates median and whiskers indicate minimum/maximum values (KPN, n = 7; KPN KF, n = 5). *\*\*P* = 0.0013, one-way Mann–Whitney U test. g) Dotplot showing neutrophils percentage in systemic blood of KPN and KPN KF mice aged to clinical endpoint (KPN, n = 8; KPN KF, n = 7). \*\*\**P* = 0.0006, one-way Mann– Whitney U test. h) Representative *Cxcl5* ISH in KPN and KPN KF tumours. Representative of five mice per genotype. Scale bar, 100 μm. i) Boxplots showing *Cxcl5* positive cells per tumour (%) of KPN and KPN KF tumours. Boxes depict interquartile range, central line indicates median and whiskers indicate minimum/maximum values (KPN, n = 6; KPN KF, n = 5). *\*P* = 0.0411, one-way Mann-Whitney U test. j) Volcano plot of differentially expressed genes in organoids derived from KPN KF tumours. k) Relative expression of *Tgfβ* ligands and chemokines from organoids derived from KPN and KPN KF tumours. KPN, n = 3; KPN KF, n = 4. l) Representative αSMA (alpha-smooth muscle actin) and Sirius Red staining of KPN and KPN KF tumours. Representative of KPN n = 7; KPN KF n = 8. Scale bar, 200 μm. m) Graphs showing Sirius Red positivity (%) and αSMA positive cells (%) of KPN and KPN KF tumours. Boxes depict interquartile range, central line indicates median and whiskers indicate minimum/maximum values (KPN n = 7; KPN KF n = 8). *\*P =* 0.0103 (SIR RED), one-way Mann–Whitney U test.

Nonetheless, we have previously shown that a critical feature of metastasis in the Notch1-driven KPN tumours is the TGFβ pathway-mediated neutrophil infiltration^32^. However, we detected a lack of systemic circulating neutrophil accumulation in the blood of KPN KF mice (Figure 6g). Given that chemokines such as *Cxcl5*, are implicated in neutrophil attraction, we assessed expression of *Cxcl5* and found significantly reduced expression in the epithelium of KPN KF tumours (Figure 6h, i). To determine how loss of wild-type *Kras* alters the TGFβ pathway and metastasis, we examined the transcriptome of tumour-derived KPN and KPN KF organoids. Interestingly, we found that NOTCH1-mediated gene expression of *Tgfb2* and chemokines, such as *Cxcl1, Cxcl3* and *Cxcl5*, was downregulated in KPN KF organoids (Figure 6j, k). Moreover, GSEA analysis showed lack of TGFβ signalling in KPN KF organoids (Extended Figure 4d). Furthermore, along with the low expression of neutrophil markers (Figure 6h, i), *Tgfb2* expression was lowered in KPN KF tumours (Extended Figure 4e). In addition, we observed clear differences in the stromal microenvironment of the KPN KF tumours, with significantly low stromal collagen deposition (picrosirius red), again, contrasted by collagen high KPN tumours (Figure 6l, m).

Given that we have previously shown that KPN tumour cells generate an immunosuppressive, pro-metastatic environment, we assessed the ability of KPN KF tumour cells to produce such an immunosuppressive environment to generate distant metastasis. To address this experimentally, we performed orthotopic intrasplenic transplantation of KPN and KPN KF tumour derived organoids in syngeneic recipient mice (Figure 7a). All mice with KPN organoid transplantation showed metastasis to either liver or lung (Figure 7b). However, the metastasis burden was significantly blunted in KPN KF organoid transplanted mice (Figure 7b, c). Remarkably, we saw a significant increase in immune cell infiltration in KPN KF transplants compared to the KPN control tumours (Figure 7d, e). Crucially, the observed impact upon immune and inflammatory infiltrate was independent of the size of metastatic deposits found in the liver – the decreased neutrophil and increased lymphocyte association with metastatic deposits in KPN KF transplants versus KPN transplants was maintained when analysis was restricted to tumours of comparable size (Extended Figure 5a, b). Moreover, the apparent impact upon immune cell infiltration was also confirmed in the KPN KF GEMM tumours (Extended Figure 4g). Together, this indicates that loss of wild-type *Kras* with epithelial NOTCH1 activation in KPN tumours activates Wnt, restricts the TGFβ mediated neutrophil recruitment required to generate an immunosuppressive, pro-metastatic niche and blunts the metastases of tumours lacking wild-type *Kras* (Figure 7f).

**Figure 7:**
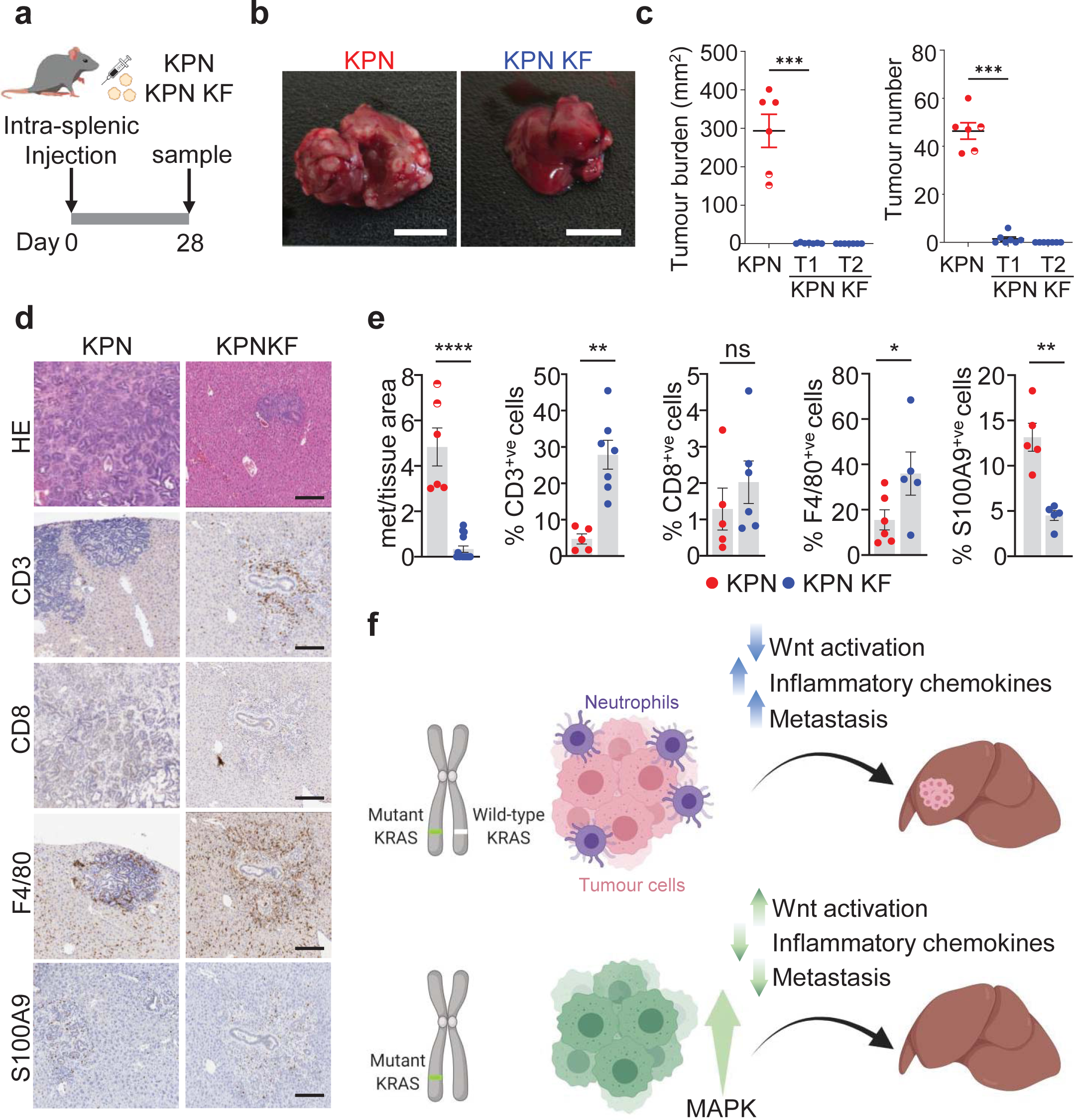
Wild-type *Kras* deficient KPN cells show reduced metastatic capacity and increased immune infiltration. a) Schematic showing the intrasplenic transplantation of KPN and KPN KF organoids. b) Images of liver tumour burden from KPN and KPN KF organoid transplant mice. Representative of six mice per organoid transplant. Scale bar, 1 cm. c) Quantifications of liver tumour burden and tumour number from KPN and two KPN KF (T1 and T2) organoid transplants. Semi circles include lung metastasis burden. *\*\*\*P* = 0.0006, one-way Mann–Whitney U test. d) Representative HE, CD3, CD8, F4/80 and S100A9 IHC in KPN and KPN KF transplant mice. Representative of six mice per organoid line. Scale bar, 200 μm. e) Quantifications from **d**. KPN KF T1 and T2 with tumour burden represented together as KPN KF. *\*\*\*\*P* = 2.5×10^-5^, *\*\*P* = 0.0013 (CD3), *\*\*P* = 0.0079 (S100A9), *\*P* = 0.02, one-way Mann–Whitney U test. f) Schematic model showing the role of wild-type KRAS in KRAS mutant CRC. Loss of wild-type *Kras* in KPN KF tumours promotes tumour initiation with WNT activation, an enhanced immune infiltrate and blunted metastasis.

## Discussion

Activating mutations in proto-oncogenes like *Kras* are the initial events in the development of cancer followed by sequential progression involving additional genetic hits in tumour suppressor genes. Here we provide evidence that loss of wild-type *Kras* facilitates and accelerates tumour initiation, but drastically alters tumour evolution in CRC. We demonstrate that loss of wild-type *Kras* increases the dependence of these tumours on the MAPK pathway thereby making them susceptible for targeting through MEK1/2 inhibition. In addition, we find that loss of tumour suppressors like p53 together with loss of wild-type *Kras* leads to faster tumour initiation but altered progression in *Kras* mutant tumours.

The role of wild-type RAS isoforms is contradictory in tumours. While the biochemical properties and transforming potential of RAS oncoproteins suggest a dominant mechanism for these mutant proteins during tumorigenesis, several studies have argued for and against the antagonizing properties of wild-type RAS in the presence of oncogenic variants^18,33,34^. There is also evidence that this is context dependent based on the RAS isoform and the tissue type under investigation^35^. Early reports suggested that presence of wild-type *Hras* or *Nras* can suppress tumourigenic phenotypes of mutant *Ras* genes^36^. While loss of wild-type Kras was shown to promote activation of all RAS isoforms in a leukaemia model^37^, loss of wild-type *Nras* did not alter the tumorigenic behaviour of mutant NrasG12D in a hematopoietic cancer model^38^. Our genetic and functional analyses using *Kras* mutant mouse models of colorectal cancer suggest that wild-type *Kras* plays a significant role in the cancer cell fitness, evolution and therapeutic susceptibility of *Kras* mutant cells *in vivo*.

Recent reports have highlighted that not all *KRAS* mutations are equal, with each *KRAS* mutation having unique biochemical, signalling and functional properties in cells^5–7^. Our data suggest that these well-characterised isoform-specific differences may be further compounded by allelic changes such as loss of the wild-type allele. Recent work from Ambrogio et al., suggested that dimerization of oncogenic KRAS is essential to sustain oncogenic function and the growth inhibitory effect of wild-type KRAS is linked to its dimerization with mutant Kras^19^. We show that wild-type *Kras* suppresses phenotypes associated with mutant *Kras* in colorectal models *in vivo*. While KRAS-driven CRC and PDAC are resistant to MEKi, the *KRAS* allelic configuration has been shown to modulate MEKi sensitivity in AML and CRC cell lines^1^. Our data show that loss of wild-type *Kras* causes cancer cells to amplify oncogenic signalling to optimize growth, leading them to be more dependent or “addicted” to certain activated pathways that become therapeutic susceptibilities. The enhanced sensitivity of *Kras*^G12D/fl^ tumours to MEKi supports this idea. Similarly, this would also suggest that in addition to screening of patients for *KRAS* mutation status, stratification of patients with *KRAS* allelic status that corresponds to selective pressure for outgrowth of tumours with increased MAPK signalling are more likely to render this subset of patients more susceptible to effector pathway inhibitors.

The idea that mutant *KRAS* gene dosage or allelic configuration can alter tumour progression has been explored recently. It is notable that as such changes are reported to be common and occur in high frequencies in *KRAS* mutant cancers, our data argue that loss of wild-type *Kras* significantly increases initiation of KRAS mutant intestinal tumours^23,26^. In contrast to *Kras*^+/G12D^ tumours, we find that *Kras*^fl/G12D^ tumours exhibit a reduction in serration, alongside reduced invasion, differentiation, and metastasis – all features previously associated with high levels of Wnt signalling in CRC. We have previously identified *Tgfb2* and chemokine secretion from the epithelium, driven by Notch activation as a key driver of metastasis in mouse models of CRC^32^. Considering the co-ordinated control of, and commonly reciprocal relationship between Wnt and Notch signalling in CRC^39^, it is intriguing that the suppression of metastasis observed in complex *Kras*^fl/G12D^ tumours occurs hand-in-hand with transcriptional enrichment of canonical Wnt-associated gene programmes. This is also consistent with reports showing that Wnt-high human CRC with elevated expression of Wnt target genes have better prognosis than Wnt-low tumours^40,41^. In the model systems reported here, Wnt-dependent transcriptional programmes were enriched in *Kras*^fl/G12D^ tumours when compared to *Kras*^+/G12D^ intestinal tumours, indicative of further crosstalk between the *Kras* and Wnt signalling pathways. When viewed in the round, these observations suggest complex interplay between the WNT, NOTCH and KRAS signalling axes in determining disease trajectory in colorectal cancer, and that allelic imbalances at the *KRAS* locus might skew this relationship, ultimately impacting prognosis and therapeutic sensitivity.

## Materials and methods

### Mouse models and experiments

All experiments were performed according to UK Home Office regulations (Project licence 70/8646), and reviewed by local ethical review committee at the University of Glasgow. Mice were genotyped by Transnetyx (Cordoba, TN, USA). Male and female C57BL/6J >20 g mice were induced with tamoxifen from 6 to 12 weeks of age. For intestinal studies, the alleles used were as follows - *VillinCre*^ER42^, *Apc*^fl43^, lox-stop-lox-*Kras*^G12D44^, *Braf*^V600E45^, *Trp53*^fl/fl46^ and *Rosa26*^N1icd/+47^. *Kras*^flox^ and *Braf*^flox^ were generated by the International Mouse Phenotyping Consortium (IMPC). *VillinCre*^ER^, *Apc*^fl^*, LSL-Kras*^G12D^ and *Kras*^flox^ mice were backcrossed for 10 generations onto C57BL/6J. Recombination by *VillinCre^ER^* was induced with one intraperitoneal (i.p.) injection of 80 mg/kg tamoxifen on day 0. Analysis of *VillinCre^ER^*-induced *Apc*^fl/fl^ *Kras*^+/G12D^ and *Apc*^fl/fl^ *Kras*^fl/G12D^ mice was at day 3 after induction. For tumourigenesis studies, *VillinCre*^ER^ *Apc*^+/fl^ (with *Kras*^+/G12D^ or *Kras*^fl/G12D^), *Kras*^+/G12D^, *Kras*^fl/G12D^, KP, KP KF, KPN and KPN KF mice were sampled when exhibiting clinical signs of ill-health indicative of substantial intestinal tumour burden (typically anaemia, hunching and/or weight loss), with overall survival recorded as the time from induction to sampling. For drug treatment studies, mice were randomly assigned to cohorts. The MEK inhibitor (AZD6244 or selumetinib) was administered in a concentration of 25 mg/kg twice daily via oral gavage in a vehicle of 0.5% HPMC and 0.1% Tween-80.

Intrasplenic transplantation injections were performed as previously described^32^ using a cell suspension of primary tumour organoids derived from C57BL/6 KPN or KPN KF mice. Organoid donor and recipient mice were strain matched. Metastatic tumour burden and tumour number were scored macroscopically upon dissection, with tumour burden recorded as cumulative area of metastatic deposit observed across all liver lobules from each individual mouse. For analysis of composition of size matched metastatic deposits, the mean area of metastatic deposit from the KPN KF cohort was calculated microscopically from a single H&E liver section per mouse. This value +/- 1 x standard deviation (σ) was used as a range of tumour area (21188.95 – 154971.4 µm^2^), which could in turn be used to select size-matched metastatic deposits from the KPN cohort for comparative analysis.

The smallest sample size was chosen that could give a significant difference, in accordance with the 3Rs. Given the robust phenotypes of the *VillinCre*^ER^ *Apc*^fl/fl^ model, and prediction that KRAS was essential, the minimum size sample size assuming no overlap in control vs. experimental is three animals.

### Immunohistochemistry (IHC) and RNAscope

Haematoxylin-and-eosin (H&E) staining was performed using standard protocols. Immunohistochemistry. IHC for BrdU (1:200, BD Biosciences #347580), β-catenin (1:50, BD Biosciences #610154), γ-H2AX (1:50, Cell Signalling Technologies #9718), phospho-ERK1/2 (1:400, Cell Signalling Technologies #9101), c-MYC (1:200 Abcham #ab32072), Lysozyme (1:300, Dako/Agilent #A099), αSMA (1:25000, Sigma-Aldrich #A2547), CD3 (1:100, Abcam #ab16669), CD4 (1:500, eBisoscience #14-9766-82), CD8 (1:500, eBisoscience #14-0808-82), and F4/80 (1:200, Abcam #ab6640) was performed on formalin-fixed intestinal sections using standard protocols. *In situ* hybridisation (ISH) (RNAscope) was performed according to the manufacturer’s protocol (Advanced Cell Diagnostics RNAscope 2.0 High Definition– Brown) for *Notum, Cxcl5* and *Tgfβ2*. Sections were counterstained with hematoxylin and coverslipped using DPX mountant (CellPath, UK). Picro Sirius Red staining technique was used to stain collagen within tissue sections as described previously^32^.

### Assaying proliferation *in vivo*

Proliferation levels were assessed by measuring BrdU incorporation. Mice were injected with 250µl of BrdU (Amersham Biosciences) 2 hours before being sacrificed. IHC staining for BrdU was then performed using an anti-BrdU antibody. For each analysis, 25 half crypts were scored from at least three mice of each genotype.

### Cell lines and crypt culture

Intestinal organoids were cultured in Matrigel and Dulbecco’s Modified Eagle Medium/Ham’s F-12 (Advanced DMEM/F-12, Gibco #12634010) supplemented with 2 mM L-glutamine (Gibco #A2916801), HEPES (Gibco #15630080) and 1% Penicillin-Streptomycin (Gibco #15070063)).

### ddPCR

Genomic DNA was extracted from snap-frozen cell pellets or mouse tissues using the QIAGEN DNeasy Blood and Tissue Kit. Reactions were performed with ddPCRSupermix and primers and probes (Bio-Rad) listed below. ddPCR was carried out according to Bio-Rad’s protocol. Droplets were generated on Bio-Rad’s QX200 with droplet generation oil, subjected to amplification (95°C 10 minutes, 94°C 30 seconds, 59°C 1 minute, repeated 40×, 98°C 10 minutes, 8°C hold), and read on Bio-Rad’s QX200 Droplet Reader running QuantaSoft software. Primer sequences are as follows: KRAS_G12D Forward: CTGCTGAAAATGACTGAGTA, Reverse: ATTAGCTGTATCGTCAAGG, Probe:TGGAGCTGATGGCGT with FAM fluorophore; KRAS_wt Forward: CTGCTGAAAATGACTGAGTA, Reverse: ATTAGCTGTATCGTCAAGG, Probe Sequence: TGGAGCTGGT-GGCG with HEX fluorophore.

### RAS activation assay

Organoids were plated two days prior the assay in Matrigel. For starvation the organoids were washed with PBS and fed ADF without EGF was for 30 min. After starvation organoids were quickly washed with ice-cold PBS, collected and disrupted prior to cell lysis. The following steps were performed using the Cytoskeleton RAS-activation assay biochem kit (#BK008) according to the manufacturer’s instructions.

### RNA Extraction, qRT-PCR and RNA-Sequencing

RNA was extracted from whole tissue, organoids or tumour tissues using QIAGEN RNAeasy kit (Cat no. 74104) according to the manufacturer’s instructions. 1 µg of total RNA was reverse transcribed to cDNA using a High-Capacity cDNA Reverse Transcription Kit (Thermo Scientific, Cat 4368814). qPCR was performed in duplicates for each biological replicate in a 20 µl reaction, with 10 µl of 2X DyNAmoHSmaster mix (Thermo Scientific, Cat no. F410L), 2 µl cDNA and 0.5mM of each of the primers. The reaction mixture without a template was run in duplicate as a control. *Gapdh* was used to normalise for differences in RNA input. For RNA sequencing, RNA integrity was analyzed using NanoChip (Agilent RNA 6000 Nano kit, 5067-1511). A total of 2 µg RNA was purified by poly(A) selection. The libraries were run on the Illumina NextSeq 500 using the High Output (75 cycles) kit (2 × 36 cycles, paired-end reads, single index). Analysis of RNA-seq data was carried out as previously described^48^. Gene set enrichment analysis was performed using GSEA version 4.1.0 software (Broad Institute).

### SDS-PAGE and Western Blotting

Organoid pellets were washed in ice-cold PBS and lysed in RIPA buffer. Protein concentration was determined using BCA protein assay (Thermo scientific # 23225). 20ug protein were separated on a 4-12% gradient pre-cast gel (Novex #NP0322PK2) in MOPS running buffer. Samples were transferred onto PVDF membrane (Millipore, #IPFL00010). Primary antibodies were diluted as indicated in 5% BSA/TBS-T (Sigma #A9647) and incubated overnight at 4°C (KRASG12D 1:1000, CST #14429; ß-actin 1:2000, SIGMA #A2228, ERK1/2 1:1000, CST #4695; pERK1/2 1:1000, CST #9101, MEK1/2 1:1000, CST ##8727 pMEK1/2 1:1000, CST #2338; AKT 1:1000, CST #9272; pAKT (Ser473) 1:2000, CST #4060, PTEN 1:1000, CST #9559). Secondary antibody (α-rabbit Dako #P0448, α-mouse Dako #P0447) was diluted 1:2000 in 5% BSA/TBS-T and incubated for 1h at RT.

### Statistics

Statistical analyses were performed using Graphpad Prism (v8) software (La Jolla, CA, USA). Statistical comparisons for survival data were performed using Mantel-Cox (Log Rank) test. Mann-Whitney-*U* tests were performed using Graphpad Prism. In each figure legend, the data and errors are shown, and the relevant statistical test is specified.

## Author contributions

A.K.N., S.K.F, A.D.C and O.J.S. designed the research. A.K.N., S.K.F., L.M.M., C.F. and N.G. performed experiments and analysed data. A.K.N., S.K.F., N.G., K.G. and W.C., generated and analysed transcriptomic data. A.K.N., S.K.F. and C.N. performed immunohistochemistry. R.A.R., E.A. and D.S. rederived *Kras*^fl^ and *Braf*^fl^ mice. J.P.M. provided expertise and provided feedback. A.K.N., S.K.F., A.D.C. and O.J.S. wrote the paper, all authors read the manuscript and provided critical comments.

## Acknowledgements

The authors are grateful to the CRUK Beatson Institute Central Services, Histology Services, Transgenics and BSU for technical support and all members of the Sansom lab for discussion of the data and the manuscript. O.J.S. was supported by CRUK grants A21139, A12481, A17196 and A31287 and ERC Starting grant 311301. A.K.N. was supported by Novartis grant awarded to O.J.S. A.K.N. was supported by Academy of Finland grant. S.K.F. was supported by Pancreatic Cancer UK Future Leaders Academy - awarded to O.J.S and J.P.M. K.G. was supported by CRUK Grand Challenge Specificancer Grand Challenge Consortium (A29055 awarded to O.J.S.). C.F, R.A.R, E.A, D.S, C.N, W.C, and A.D.C. were supported by CRUK Beatson Institute core funding A17196, A31287 – awarded to O.J.S. J.P.M was supported by CRUK Beatson Institute core funding A17196, A31287 and A29996.

---

**Extended Figure 1:**
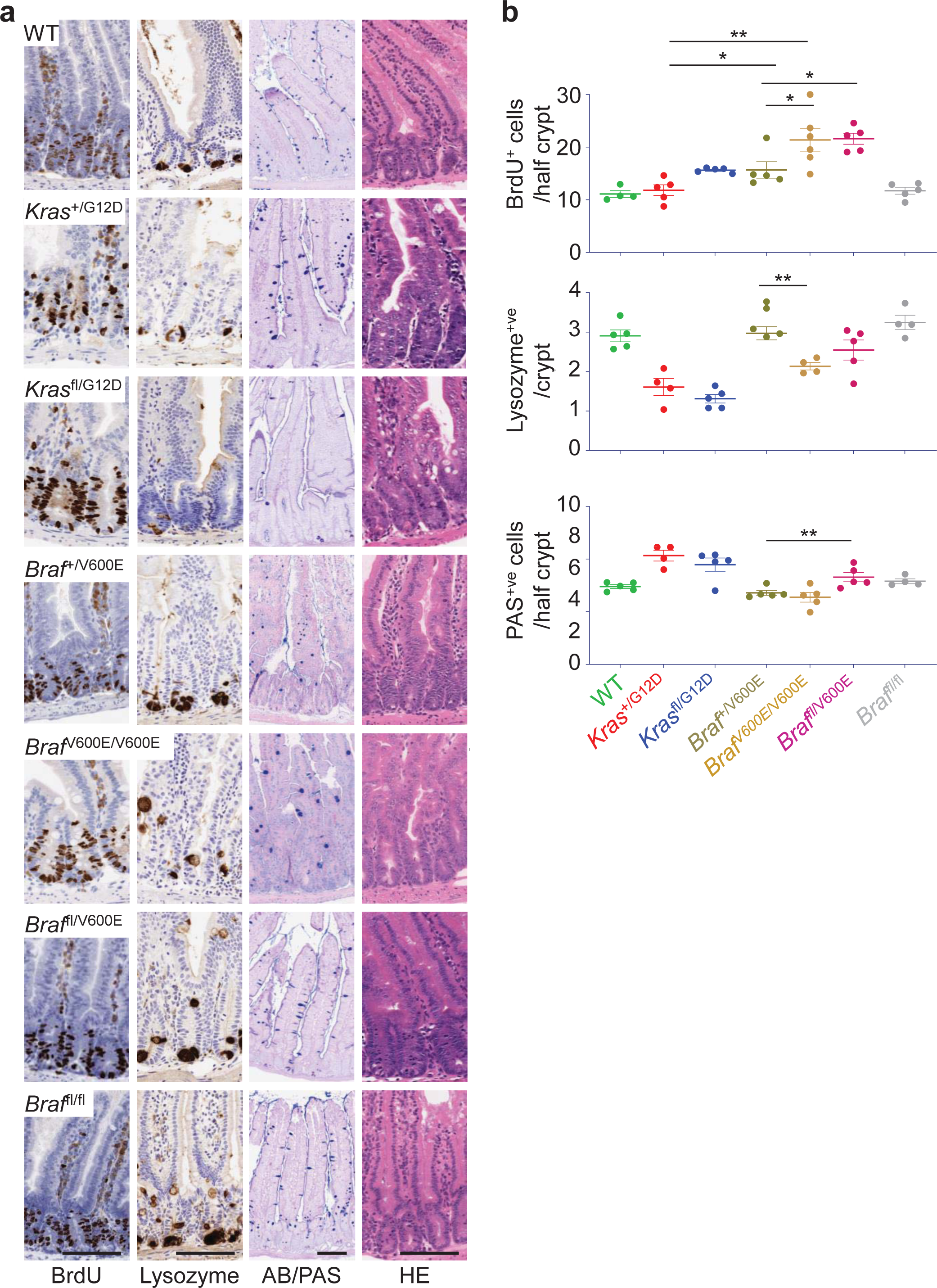
*Braf*^V600E^ gene dosage change alters intestinal homeostasis. a) Representative BrdU, Lysozyme, AB/PAS and H&E IHC of *Kras*^+/+^, *Kras*^+/G12D^, *Kras*^fl/G12D^, *Braf*^+/V600E^, *Braf*^V600E/V600E^, *Braf*^fl/V600E^ and *Braf*^fl/fl^ mouse small intestine, sampled 4 days post Cre-induction. Scale bar, 100 μm. b) Top: Quantification of the number of BrdU-positive cells from at least 25 half-crypts in SIfrom (a). Data are mean ± s.e.m, (*Kras*^+/+^, n = 4; *Kras*^+/G12D^, n = 5; *Kras*^fl/G12D^, n = 5; *Braf*^+/V600E^, n = 5; Braf^V600E/V600E^, n = 6; *Braf*^fl/V600E^, n = 5; *Braf*^fl/fl^, n = 5), \*\**P* = 0.0079. Middle: Number of Lysozyme-positive cells from at least 25 crypts in SIof *Kras*^+/+^, *_Kras_*+/G12D_, *Kras*_fl/G12D_, *Braf*_+/V600E_, *Braf*_V600E/V600E_, *Braf*_fl/V600E _and *Braf*_fl/fl _in SI from (d)_ Data are mean ± s.e.m, (*Kras*^+/+^, n = 5; *Kras*^+/G12D^; n = 4; *Kras*^fl/G12D^, n = 5; *Braf*^+/V600E^, n = 5; Braf^V600E/V600E^, n = 4; *Braf*^fl/V600E^,n = 5; *Braf*^fl/fl^,n = 4), \*\**P* = 0.0028. Bottom: Number of PAS-positive cells from at least 25 half-crypts in SI. *Kras*^+/+^, *_Kras_*+/G12D_, *Kras*_fl/G12D_, *Braf*_+/V600E_, *Braf*_V600E/V600E_, *Braf*_fl/V600E _and *Braf*_fl/fl _in SI from (d)_ Data are mean ± s.e.m, (*Kras*^+/+^, n = 5; *Kras*^+/G12D^, n = 4; *Kras*^fl/G12D^, n = 5; *Braf*^+/V600E^, n = 5; Braf^V600E/V600E^, n = 5; *Braf*^fl/V600E^, n = 5; *Braf*^fl/fl^, n = 4). \*\**P* = 0.0079, All P values generated using one-way Mann–Whitney U test.

**Extended Figure 2:**
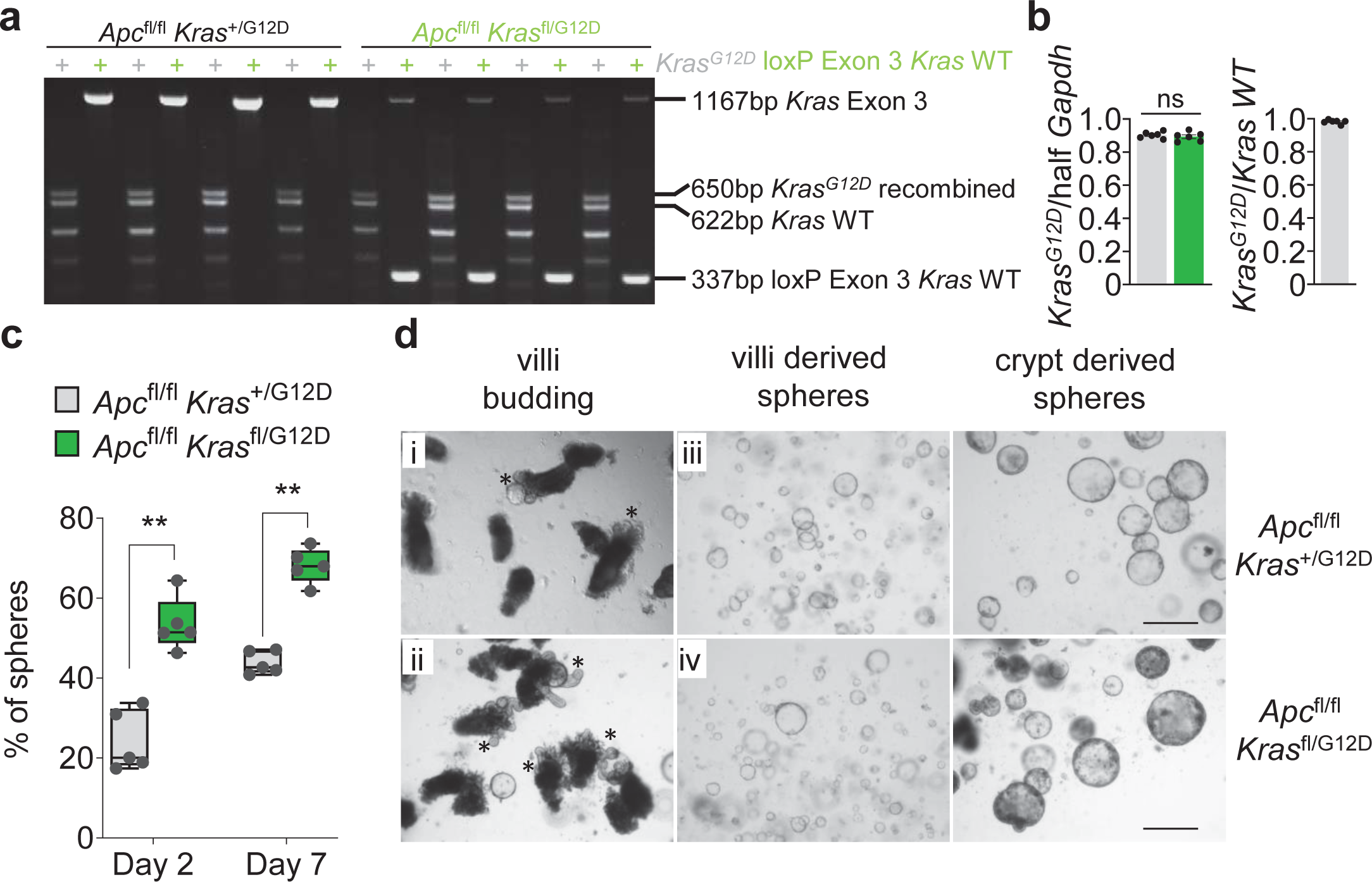
Increased de-differentiation capacity of *Apc Kras*^G12D^ villi lacking wild-type *Kras*. a) Representative data of PCR analysis of *Kras* loci in *Apc*^fl/fl^ *Kras*^G12D/+^ and *Apc*^fl/fl^ *Kras*^fl/G12D^ organoids showing the recombined *Kras*^G12D^ and *Kras*^fl^ (*Kras* exon 3 deletion). b) Bar graph showing KrasG12D copy number in *Apc*^fl/fl^ *Kras*^G12D/+^ and *Apc*^fl/fl^ *Kras*^fl/G12D^ organoids, expressed relative to half of GAPDH. In most of the cases analyzed, the Kras showed no mutant amplification. *Apc*^fl/fl^ *Kras*^G12D/+^; *Apc*^fl/fl^ *Kras*^fl/G12D^ n = 6, *ns*=not significant, one-way Mann–Whitney U test. KrasG12D relative to wild-type in *Apc*^fl/fl^ *Kras*^G12D/+^ showing Kras allelic balance. c) Boxplots showing percentage of spheres forming from *Apc*^fl/fl^ *Kras*^+/G12D^ and *Apc*^fl/fl^ *Kras*^fl/G12D^ mouse intestinal-derived organoids at day 2 and day 7 after seeding. Boxes depict interquartile range, central line indicates median and whiskers indicate minimum/maximum values, representative of 5 biological replicates of organoids generated from 5 individual mice per genotype. \*\**P* = 0.004 (day2, day7) one-way Mann–Whitney U test. d) Representative images of *Apc*^fl/fl^ *Kras*^+/G12D^ and *Apc*^fl/fl^ *Kras*^fl/G12D^ villi and crypt derived organoid spheres. Representative of 5 biological replicates. Asterisks marks budding spheres.

**Extended Figure 3:**
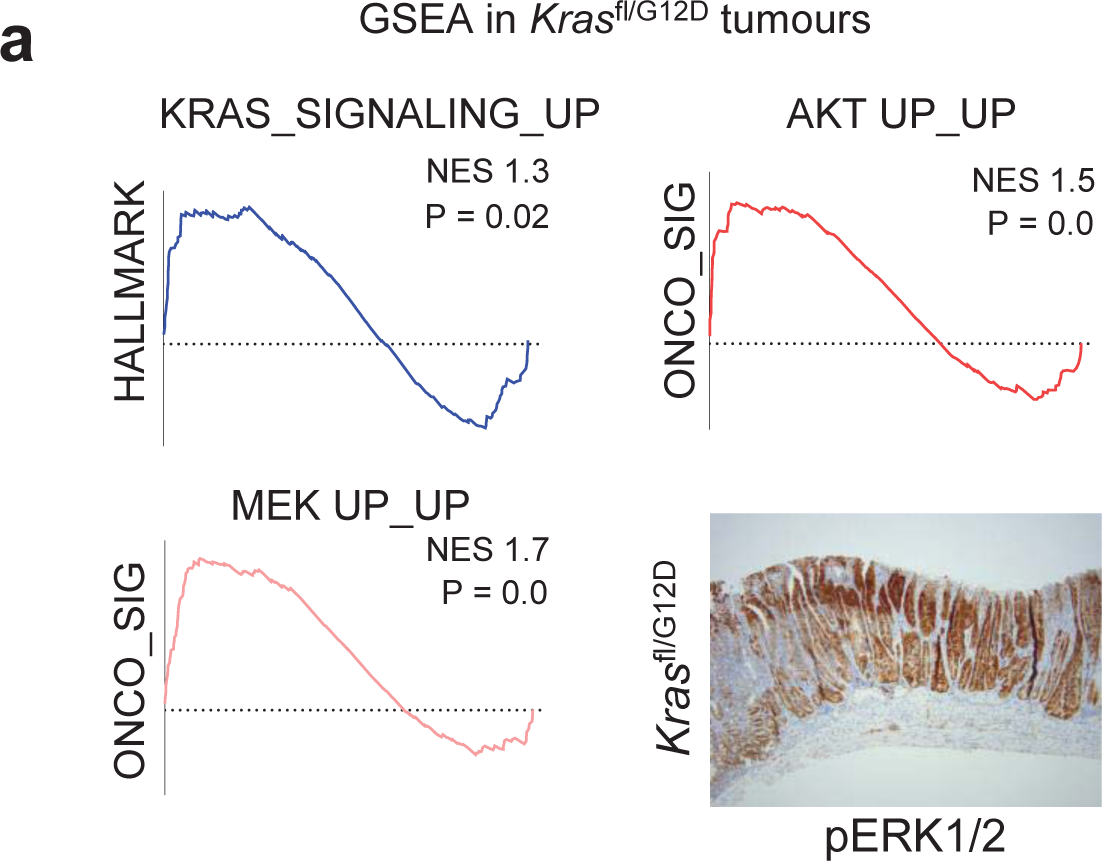
Loss of wild-type *Kras* in Kras^G12D^ mutant intestinal tumours leads to transcriptional enrichment of effector pathways. a) GSEA with the KEGG, Hallmark and oncogenic signatures collection yields an array of highly significant enriched gene sets in *Villin*^CreER^ *Kras*^fl/G12D^ tumours. Selection of enrichment plots representing *KRAS*, *MEK* and *AKT* signalling.

**Extended Figure 4:**
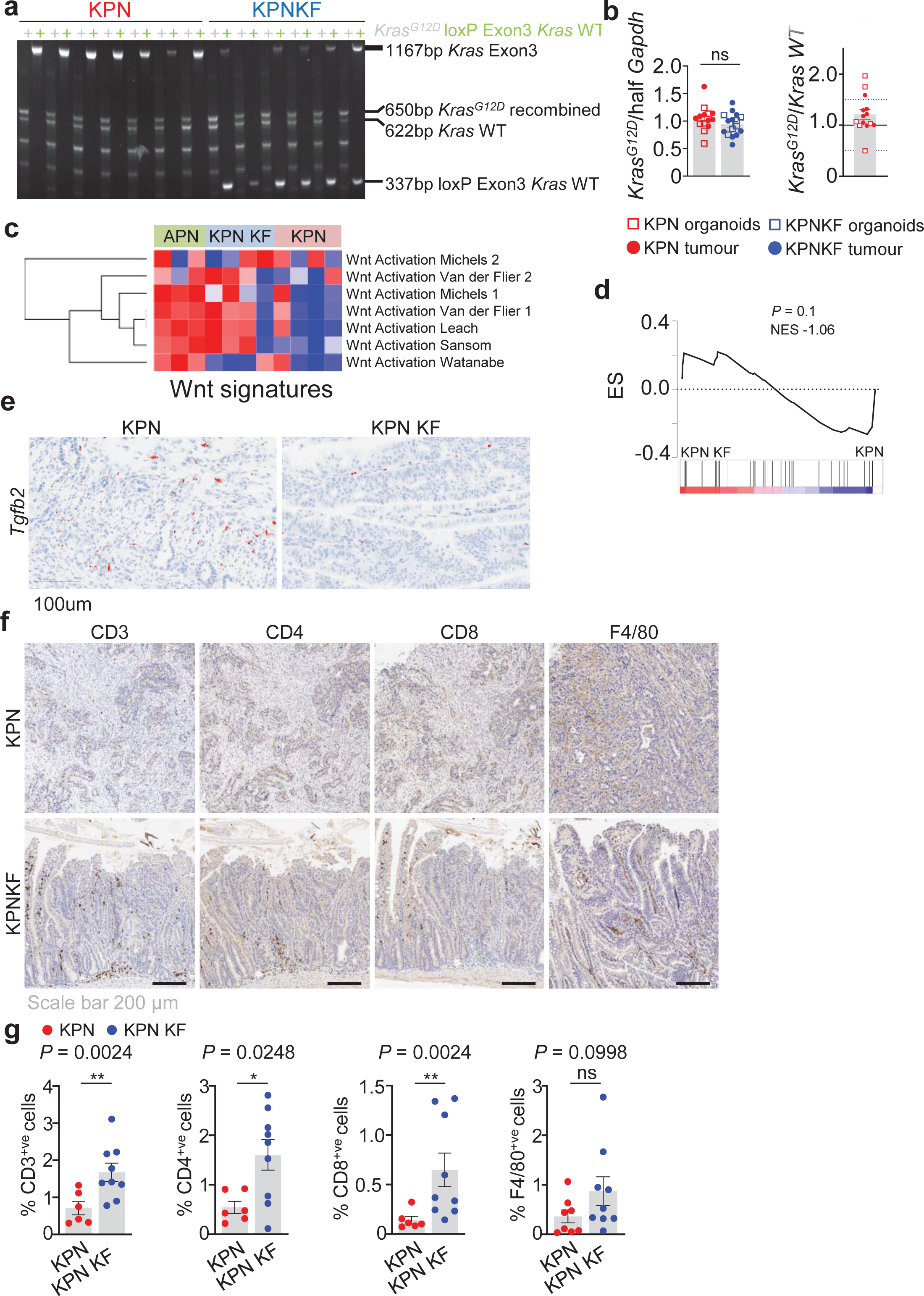
Wild-type *Kras* deletion promotes WNT activation and alters TGF*β* levels in KPN tumours. a) Representative data of PCR analysis of *Kras* loci in KPN and KPN KF organoids showing the recombined *Kras*^G12D^ and *Kras*^fl^ (*Kras* exon 3 deletion). b) Bar graph showing KrasG12D copy number in KPN and KPN KF organoids and tumours, expressed relative to half of GAPDH. In most of the cases analyzed, the Kras showed no mutant amplification. KPN n = 14; KPN KF n = 16, *P*=0.07, one-way Mann– Whitney U test. KrasG12D relative to wild-type Kras showing most KPN samples (n = 14) have allelic balance and two organoid lines with a mutant Kras amplification. c) Heatmap of WNT Activation signatures for *Apc*^+/fl^ *Trp53*^fl/fl^ *Rosa26*^N1icd/+^ (APN), *Kras*^fl/G12D^ *Trp53*^fl/fl^ *Rosa26*^N1icd/+^ (KPN KF) and *Kras*^+/G12D^ *Trp53*^fl/fl^ *Rosa26*^N1icd/+^ (KPN) tumours. d) GSEA of hallmark *Tgfβ* signalling (GSE15871) in KPN KF tumour organoids compared to KPN. NES = normalised enrichment score. e) Representative *Tgfβ2* ISH in KPN and KPN KF tumours. Representative of five mice per genotype. Scale bar, 100 μm. f) Representative CD3, CD4, CD8 and F4/80 IHC in KPN and KPN KF tumours. Representative of six mice per genotype. Scale bar, 200 μm. g) Bar graph showing the quantifications from **e** of KPN and KPN KF tumours. *P* values indicate, one-way Mann–Whitney U test.

**Extended Figure 5:**
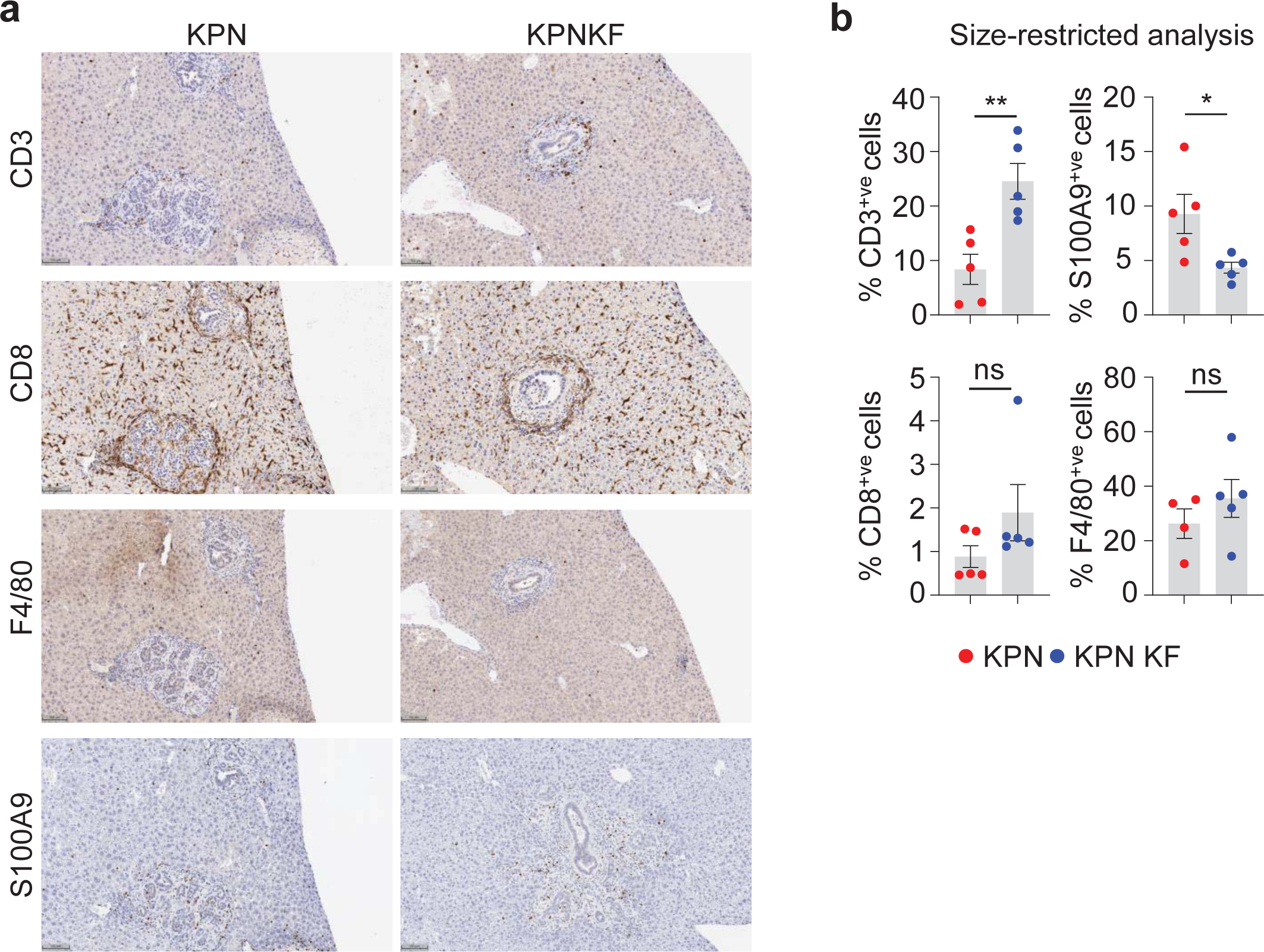
Wild-type *Kras* deficient KPN cells show increased immune infiltration also in size matched tumours. a) Representative CD3, CD8, F4/80 and S100A9 IHC in KPN and KPN KF transplant mice. Representative of six mice per organoid line. Scale bar, 100 μm. b) Bar graph showing the quantifications from **a** of KPN and KPN KF tumours. *\*\*P* = 0.0079, *\*P* = 0.0159, ns = non significant, one-way Mann–Whitney U test.

## Notes

### Competing Interest Statement

The authors have declared no competing interest.

